# Addressing biases in gene-set enrichment analysis: a case study of Alzheimer’s Disease

**DOI:** 10.1101/2023.08.13.553133

**Authors:** Artemy Bakulin, Noam B Teyssier, Martin Kampmann, Matvei Khoroshkin, Hani Goodarzi

## Abstract

Inferring the driving regulatory programs from comparative analysis of gene expression data is a cornerstone of systems biology. Many computational frameworks were developed to address this problem, including our iPAGE (**i**nformation-theoretic **P**athway **A**nalysis of **G**ene **E**xpression) toolset that uses information theory to detect non-random patterns of expression associated with given pathways or regulons^1^. Our recent observations, however, indicate that existing approaches are susceptible to the biases and artifacts that are inherent to most real world annotations. To address this, we have extended our information-theoretic framework to account for specific biases in biological networks using the concept of conditional information. This novel implementation, called pyPAGE, provides an unbiased way for the estimation of the activity of transcriptional and post-transcriptional regulons.

To showcase pyPAGE, we performed a comprehensive analysis of regulatory perturbations that underlie the molecular etiology of Alzheimer’s disease (AD). pyPAGE successfully recapitulated several known AD-associated gene expression programs. We also discovered several additional regulons whose differential activity is significantly associated with AD. We further explored how these regulators relate to pathological processes in AD through cell-type specific analysis of single cell gene expression datasets.

## INTRODUCTION

Over the past two decades, gene expression analysis has emerged as an effective approach for defining cell states at a molecular level. For example, disease signature genes, defined as genes that are up or down-regulated in diseased cells, have long been used as a proxy for pathological cell identity^2^. However, even in the early days, it was clear that for complex diseases like cancer, signature genes fail to reproduce in independent studies^3^. This was not surprising, as it is the collective disruptions in cellular pathways and processes, rather than dysregulated expression of individual genes themselves, that underlies cellular pathologies. This understanding gave rise to a suite of computational analyses that perform pathway-level or gene set-level comparisons between samples. Similarly, for previously annotated regulatory networks, such as transcription factor regulons, the target gene-sets were often used to assess the differential activity of master regulators across conditions^4^. These target regulons were either annotated using biological data (such as ChIP-seq^5^ and CLIP-seq^6^) or based on sequence binding preferences (AlignACE^7,8^, FIRE^9^, DeepBind^10^, DeepSEA^11^, DanQ^12^). We have previously shown that application of this methodology to large scale gene expression datasets reveals the underlying regulatory programs that are key to the emergence or maintenance of a given cellular state^1^. However, as we will discuss here, additional challenges needed to be addressed to further improve our ability to detect perturbations in regulatory programs.

There are myriad tools implementing pathway level analysis; Garcia-Campos et al^13^ categorize these tools into two main categories: over-representation analysis (ORA) methods and Functional Class Scoring (FCS) methods. Over-representation analysis on the concept that the top-scoring genes should carry the highest weight; therefore, the ORA methods operate under the null hypothesis that if a number of differentially expressed genes is chosen, the fraction of genes that belong to any given pathway does not exceed the fraction expected by chance. In contrast, FCS methods operate under the assumption that pathways might consist of many genes that show heterogeneous yet coordinated changes in expression. Such methods make use of all available measurements and often do not rely on arbitrary thresholding.

In these methodologies, pathway annotation databases, such as Gene Ontology^14,15^ and MSigDB^3,16^, can be used to represent pathways as gene sets or binary vectors where a gene can either belong to a given pathway, or not. However, this commonly used representation of pathway membership creates an intrinsic bias for several reasons. First, genes are multifunctional and some genes are far more studied than others and therefore are simultaneously annotated as part of many pathways. Second, the more abundant a given protein, the more often it is detected in various assays (such as affinity purification); therefore, abundant proteins similarly tend to be categorized as members of many complexes and gene sets. Such biases can have a broad impact on target selection and pathway identification in large-scale datasets. For example, the observation that a protein that is only annotated as part of a few gene sets is differentially expressed narrows down the selection of potentially relevant pathways much more than a similar observation about a protein that is present in many gene sets. To our knowledge, no available pathway-analytical tool explicitly takes such biases into account.

Here, we propose a pathway analytical approach that explicitly models the representation bias present in gene-set annotations, and uses an underlying deterministic statistic rather than permutation-based approaches. This framework is an extension of our iPAGE package developed more than a decade ago^1^. In iPAGEv1.0 we used mutual information to directly estimate the dependency between expression values and pathway annotations. In contrast, pyPAGE relies on calculating conditional mutual information, which allows for explicit consideration of annotation bias as a covariate. Here, we provide a number of benchmarks showcasing the utility of pyPAGE as a robust pathway analytical tool that surpasses its predecessors. And as we will demonstrate, pyPAGE can effectively reveal the regulatory mechanisms underlying biological observations, which is a crucial first step towards a molecular understanding of their origin and maintenance.

The generalizability of pyPAGE allows for its application to a variety of different data modalities in order to infer regulatory changes at various molecular levels. We took advantage of this to carry out a comprehensive analysis of gene expression programs in Alzheimer’s disease. While AD is currently characterized at molecular, histological, and symptom levels^17^, the regulatory networks, both transcriptional and post-transcriptional, that accompany and likely drive cellular pathogenesis in AD are not fully explored. Leveraging pyPAGE, we have generated novel insights into the role of key master regulators, including transcription factors (TF), RNA binding proteins (RBP), and microRNAs (miRNAs), in shaping the pathological transcriptome in AD. First, we describe deregulation patterns of their regulons based on differential gene expression and differential transcript stability measurements inferred from MSBB RNA sequencing data^18^. Next we provide a cell-type specific characterization of differential activity of the identified regulons using two single-cell RNA sequencing datasets^19,20^. We also leverage these datasets to uncover regulons whose differential activity patterns remained hidden in the bulk data. Finally, we demonstrate the utility of pyPAGE to perform stratification of patients based on the inferred activity of the regulons.

## RESULTS

### Gene-set annotations often form scale-free networks

In every annotation of biological regulons, the number of gene sets a gene is assigned to varies significantly among all genes. Typically, this measure of gene-set membership reflects the number of functional interactions a gene engages in. Current analytical tools assume gene-set membership to follow a normal distribution (**Figure 1A**). However, in practice this assumption rarely holds true and gene-set membership instead follows the power law (**Figure 1A**). We have demonstrated this across multiple commonly used gene-set annotations: molecular pathways, transcription factor target genes, RNA-binding protein regulons and miRNAs targets (**Figure S1, A-D**). In each of these annotations, the degree distribution of the gene-set membership could be closely approximated using the power law function (R^2^ > 0.9), which typically peaks at 1 with a long distribution tail. This implies that like most biological networks, gene-set annotations also form scale-free networks.

**Figure 1.**
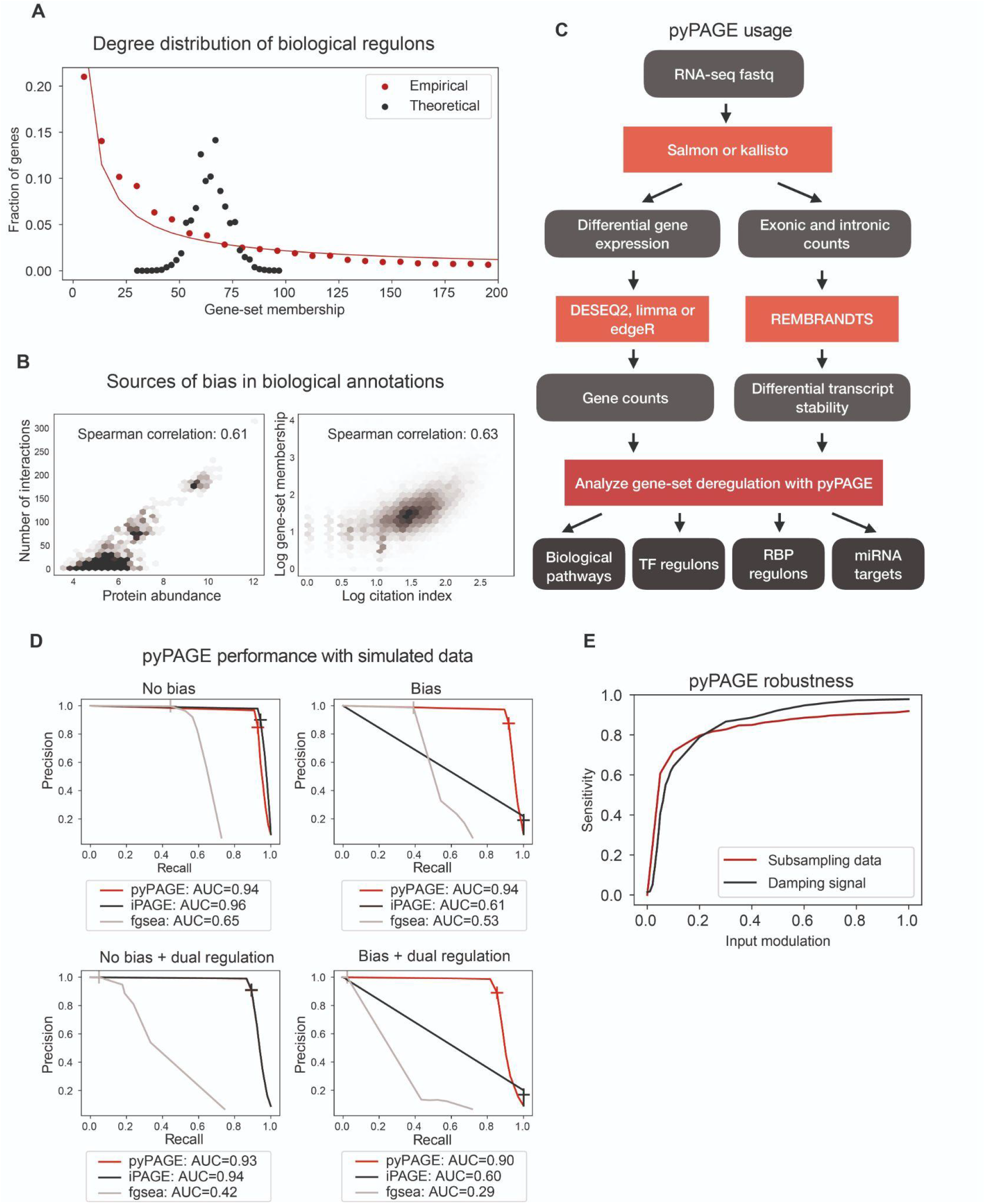
pyPAGE is a novel framework for inference of differentially regulated gene-sets. (A) Comparison between the theoretical and empirical degree distributions of gene-set membership in gene-set annotations. The red line represents the curve-fitting of the power law function to the observed distribution. **(B)** Scatter plots representing major sources of bias in biological annotations. The left panel represents the association between the protein abundance and the number of interactions a gene has in the STRING database. The right panel represents the association between the citation index of a particular gene and its gene-set membership in an annotation of biological pathways. For each association, we also report correlation between values. **(C)** Schematic of the pipeline we propose for the analysis of bulk RNA-seq data using pyPAGE. **(D)** Precision-recall curves demonstrating the performance of pyPAGE and benchmarking it against iPAGE and fgsea. The analysis was made in 4 simulated scenarios with and without added biases and with or without dual regulation patterns. As a general metric of performance we report PR-AUC score, also cross glyphs mark the performance at p-value threshold equal to 0.01. **(E)** A plot demonstrating robustness of pyPAGE to changes in quality of the input data. Two curves correspond to modulation of the size of the input data and the intensity of the input signal.

Further analysis of these gene-set membership distributions revealed that in many cases the shape of the right tails significantly deviates from the power law (**Figure S1A and D**). Such deviations indicate a bias in the underlying annotations. We posit that this bias arises from the experimental and analytical techniques that are used to compile them. For example, we observed that the more a given gene is studied the more likely it is to have multiple entries in a knowledge-based annotation (**Figure 1B, right panel**). Another source of bias may be technical limitations and low sensitivity of detecting weak functional interactions. Consistently, we have observed that in human interactome the number of physical interactions for a given protein correlates with this protein abundance (**Figure 1B, left panel**). Overall, these findings indicate that there are important structural biases in the commonly used gene-set annotations. As we will demonstrate below, these inherent biases, which are by and large ignored in gene-set enrichment analysis, have an impact on the insights that can be gleaned from the underlying data using current methodologies.

### pyPAGE, an unbiased approach for gene-set enrichment analysis

Current gene-set enrichment analysis tools assume that every gene in a regulon is equally informative of regulon activity. However, as we demonstrated above, gene-set membership significantly varies between genes. As a consequence, differential expression of genes with specific functions provides stronger evidence for deregulation of its corresponding gene-sets than changes in expression of a more promiscuous gene. Here, we describe pyPAGE, an improved gene-set enrichment analysis method that directly addresses biases and imbalances in annotations. pyPAGE is an extension of our previously published iPAGE framework^1^ and can be used to analyze various gene-set annotations such as Gene Ontology, MSigDB and regulons of TFs, RBPs and microRNAs. To correct for imbalances in gene-set membership, pyPAGE uses the concept of conditional mutual information^21^ and measures the dependency between input data (e.g. gene expression) and a specific gene-set while taking into account the total number of gene-sets every given gene belongs to. The statistical significance of this dependency is then determined using a permutation test. Like its predecessor, pyPAGE detects any non-random patterns of enrichment or depletion of target genes and even non-monotonic complex relationships are captured (e.g. dual regulators). It also does not depend on the particular distribution of the input data and can be applied to both discrete (e.g. cluster indices) and continuous (e.g. log-fold changes) inputs.

pyPAGE allows for the comprehensive analysis of gene expression changes at multiple levels, including biological pathways as well as transcriptional and post-transcriptional regulons. To standardize the analysis of biological datasets using pyPAGE, we propose the workflow described in **Figure 1C**. We start with initial preprocessing of RNA-seq data followed by estimation of differential gene expression using any of the available tools, such as DESeq2^22^, limma^23^ or edgeR^24^. Next, we apply pyPAGE to identify biological pathways and transcriptional factor regulons which are significantly associated with the differential gene expression. To analyze activity of post-transcriptional regulons one must take a step back to get the unbiased estimates of transcripts stability. Our recently introduced tool REMBRANDTS leverages comparison of intronic and exonic reads to estimate changes in RNA stability^25^. Based on these differential RNA stability measurements, pyPAGE is able to infer the activity of post-transcriptional regulons, controlled by RBPs and microRNAs. This outline describes bulk analysis. Alternatively, to better capture cell-type specific patterns of regulation, pyPAGE can be applied to single-cell measurements.

### pyPAGE outperforms existing tools on simulated datasets

As described above, gene-set annotations are often imbalanced and biased in ways that are not captured and corrected for by existing gene-set enrichment analysis tools. To assess the ability of pyPAGE to account for these biases while maintaining a high baseline performance relative to its predecessors, we used several simulation-based validation strategies. By controlling the “ground truth” signal in simulated data and setting *a priori* expectations for gene-set modulations, we were able to measure the sensitivity of pyPAGE and compare its performance to other similar tools.

The main objectives of these simulations were (i) to demonstrate that implicit biases in gene-set annotations can negatively impact the quality of gene-set enrichment analysis, and (ii) to assess whether pyPAGE effectively addresses this issue and robustly identifies deregulation of gene-sets from biased annotations. . Our secondary goal was to demonstrate that pyPAGE can detect any non-random patterns in data, both monotonic and non-monotonic. Though activity of most factors is associated with either upregulation or downregulation of their target regulons, there are nevertheless factors that function as dual regulators simultaneously exerting both effects.

Therefore, we simulated two types of datasets with monotonic and dual regulation patterns. To compare against pyPAGE, we also used its predecessor iPAGE^1^ and another commonly used tool for gene-set enrichment analysis fgsea^26^. The performance of each method was measured based on its ability to recover the artificially perturbed gene sets in each simulated dataset. We employed four simulation scenarios showcasing how well each algorithm handles bias in annotations and captures monotonic as well as non-monotonic regulatory patterns. In each case, we report relevant performance metrics (**Figure S2**). However, since the number of unperturbed gene sets in the simulation is much greater than the number of perturbed gene sets, we specifically focus on the precision-recall curves (**Figure 1D**). In these benchmarks, pyPAGE and iPAGE performed equally well predicting deregulation of gene sets from unbiased annotations (PR-AUC>0.90). However, when using biased annotations, performance of iPAGE decreased dramatically by more than 30% as indicated by changes in PR-AUC score. Performance of pyPAGE, on the other hand, remained largely unchanged. Fgsea performed poorly using the same simulated data across all tasks. Firstly, it failed to accept several gene-sets as input and could not analyze their deregulation because of imbalances in their input distributions. Secondly, it largely failed to capture non-monotonic and bimodal signals. Finally annotational biases caused its performance to fall by more than 15%. These results demonstrate that based on the criteria formulated by our two validation objectives, pyPAGE outperforms other tools in being robust to biases in gene-set annotations and identifying complex patterns in data.

### pyPAGE is both robust and sensitive in detecting dysregulated gene sets

To further assess the robustness and sensitivity of pyPAGE, we implemented several additional simulation strategies. Our goal was to demonstrate that the identified gene sets remained consistent across independent runs, even as signal-to-noise was reduced. To suppress the signal in the input data, we employed two approaches. First, we directly controlled the magnitude of the simulated gene expression changes by tuning it as a hyperparameter. In the second approach we simulated new datasets by randomly selecting and changing a subset of genes in the target gene-set. The former strategy is designed to assess the sensitivity of pyPAGE to the strength of the input signal and the latter will allow us to evaluate its robustness to incomplete data or imperfect gene-set annotations^9^. pyPAGE proved to be highly robust to the choice of simulation parameters controlling signal expressiveness within a wide range (**Figure 1E**), which is showcased by its ability to capture more than 80% of the target gene-sets even after an 80% decline in signal strength. Similarly, even after sub-sampling the genes to as low as 20% of the original counts, pyPAGE was still able to recover more than 80% of the perturbed target gene-sets. This test was especially important, since we aim to extend usage of pyPAGE to single-cell RNA-seq datasets, where substantially fewer genes are captured relative to bulk data.

### Comprehensive Analysis of Regulatory Perturbations in Alzheimer’s disease

As we showed above, pyPAGE provides a sensitive and robust approach for identifying the regulatory programs that drive changes in gene expression. Extending our work to real-world data, we sought to use pyPAGE to study regulatory perturbations in Alzheimer’s disease (AD). This disease represents a highly complex phenotype and involves dysregulation of many cellular pathways and processes and the recent explosion in the amount of transcriptomic data in AD patients presents an opportunity to systematically investigate the transcriptional and post-transcriptional regulatory programs that are affected by this disease.

Alzheimer’s disease primarily causes the atrophy of the large cortical, subcortical and a number of other areas of the brain^27^. Pathologically, it is characterized by neuronal loss and destruction of synaptic connections, which are linked to the accumulation of amyloid beta and tau protein. Previous research also points to a variety of other pathological mechanisms that may be involved in AD, such as immune dyshomeostasis^28^, oxidative stress^29^ and metal imbalance^30^. A handful of studies have sought to identify master regulators of these changes based on their own differential gene expression^31–33^. However, since the activity of proteins is often controlled at the post-translational level^34^, it cannot be reliably inferred from their mRNA expression levels alone. Therefore, we sought to instead infer their differential activity from changes in the expression of their downstream target genes. In our earlier work, we demonstrated that an information-theoretic approach can be effectively employed to capture regulatory programs that underlie complex human diseases^1^. Similarly, pyPAGE uses data from known and putative interactions between regulators and genes, allowing us to systematically infer the associations between the AD pathology and deregulation of TFs, miRNAs and RBPs. The major advantage of using pyPAGE here is that it effectively accounts for uncertainty around putative regulatory interactions and non-specific and artifactual hits, therefore improving our ability to capture real biological signals in the data.

Using publicly available RNA-seq data from the MSBB study^18^, we performed a comparison of samples from the BM36 brain region between individuals with an AD diagnosis and without, while controlling for biological variables such as sex. To capture differences in transcriptional control, we used our previously curated annotation of *cis*-regulatory elements and transcription factor binding sites^35^. Similarly, post-transcriptional regulatory changes were predicted based on the combined annotation of known RBP binding sites from various sources (see Methods) and miRNA recognition elements^36,37^. To ensure that the identified regulatory programs reflect AD pathogenesis and not changes in cellular composition artificially read out in bulk data, we also analyzed their cell-type specific activation patterns using single cell measurements, as well as their clinical relevance in predicting long-term survival of AD patients.

### Transcriptional regulatory modules associated with Alzheimer’s disease

By performing the above-described analysis of transcriptional deregulation in AD, we observed significant associations for 15 transcription factors (TFs) (**Figure 2A**). We demonstrate that changes in expression of these TF regulons occur in both female and male donors (**Figure S3A**) and in each age group (**Figure S3B**). For a number of these regulons, we also observed concomitant changes in the expression levels of their associated TFs as well (**Figure 2B**). While the correlated expression of a TF and its regulon strengthens the evidence for the observed associations, as mentioned earlier, differential expression of a TF is not the sole source of its differential activity. For example, our analysis correctly identified two well known transcriptional regulators of AD, namely KDM5A^38^ and ATF4^39^. However, only the expression of KDM5A showed a high correlation with the mean expression of its regulon (R^2^=0.85) (**Figure 2C**); whereas ATF4 demonstrates lower correlation coefficient (R^2^=0.40) (**Figure 2D**). The latter is not coincidental since ATF4 levels are known to be controlled at the level of translation^40^.

**Figure 2.**
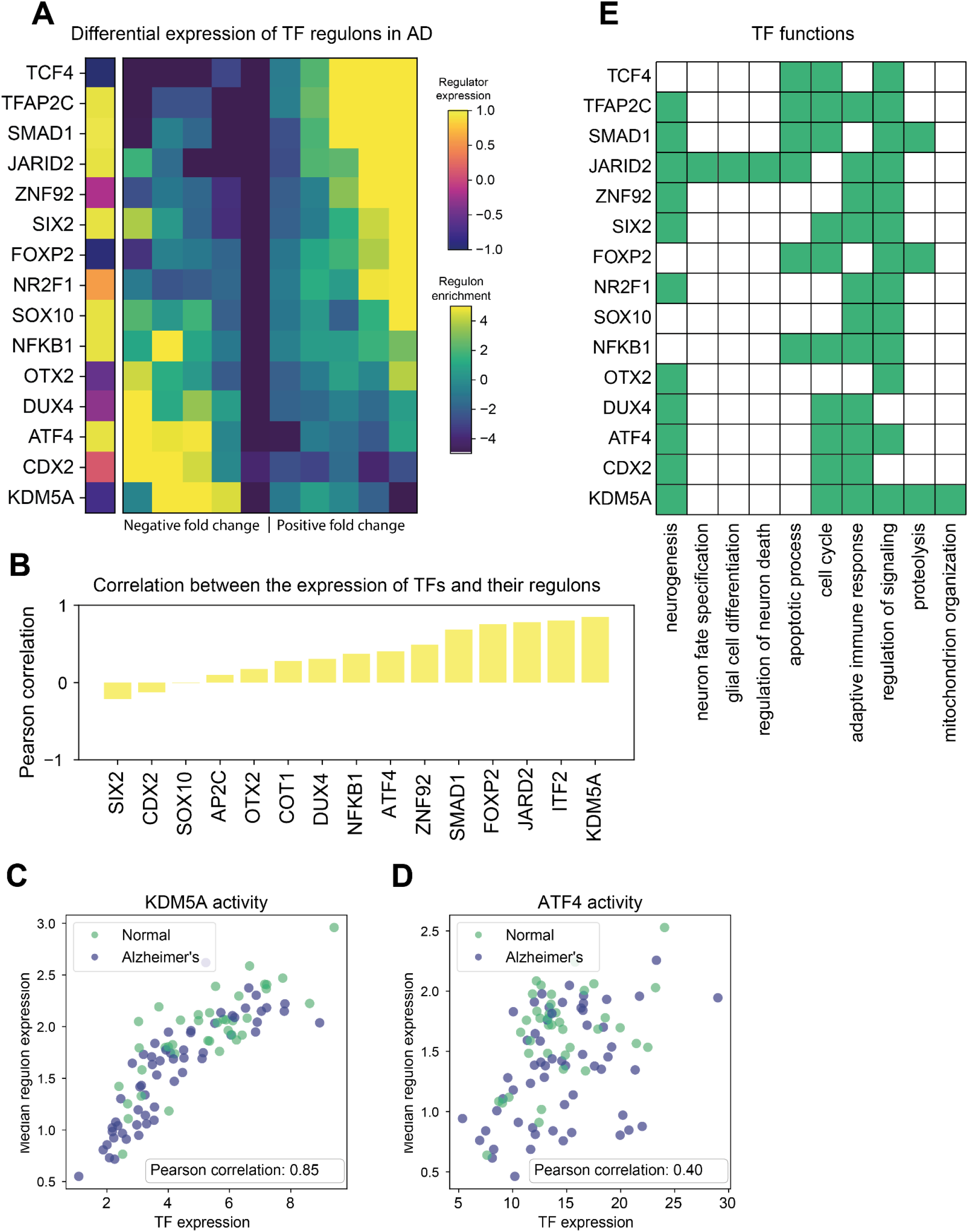
Transcription factors associated with gene expression changes in Alzheimer’s Disease. (A) Regulons of TFs differentially expressed between AD and non-AD samples discovered by pyPAGE. In this representation the rows correspond to TFs and columns to gene bins of equal size ordered by differential expression, the cells are colored according to the enrichment of genes from regulons in a corresponding bin. The leftmost column of the heatmap depicts the differential expression of the regulator itself. **(B)** Enrichment of biological pathway members among target genes of the identified TFs. In this heatmap colored cells represent significant (p-value<0.05) associations. **(C)** The barplot representing Pearson correlations between the expression of TFs and of their regulons, as measured by median TPM of its members. **(D)** The scatter plot demonstrating association between the expression of the well known AD regulator KDM5A with the expression of its regulon. **(E)** The association between the expression of another AD regulator ATF4 with the expression of its regulon.

Consistent with their implied role in AD, many of the identified TFs are key regulators of brain development and homeostasis. Systematic enrichment analysis of genes from various biological pathways within regulons of these TFs, revealed that 11 out of 15 TFs regulate molecular programs related to neurogenesis (**Figure 2E, Figure S4 and Table S2**). For example, ATF4 maintains excitatory properties of neurons^32^, whereas the generation of oligodendrocytes^42^ is controlled by SOX10. While the majority of the identified TFs function as transcriptional activators, two of them, namely JARID2 and KDM5A, are also involved in epigenetic reprogramming, which is a hallmark of AD pathology^43,44^. Additionally, several of the identified factors control adaptive immune response and protein metabolism, linking these regulons to other key hallmarks of AD: inflammation, amyloid accumulation and abnormal phosphorylation of tau protein.

### Cell-type specific patterns of deregulation for AD-associated TF regulons

Since the analysis of bulk data captures a mixture of cell types, it often requires major shifts in gene regulation. Moreover, some of the patterns observed in bulk may reflect changes in cell-type abundance rather than bona-fide reprogramming of gene expression. To account for this possibility and improve resolution of our analysis, we applied pyPAGE to the ROSMAP single-cell RNA-seq study^19^, which includes the major brain cell-types in samples from both AD and healthy donors (**Figure 3A**). For each cell-type, we estimated differential gene expression between two groups of samples and then applied pyPAGE to predict deregulated TF activity. Since the approach implemented by pyPAGE is independent of any particular distribution of the input data, it was seamlessly extended to the scRNA-seq data domain. The benefits of using pyPAGE over other tools developed for analysis of scRNA-seq data (e.g., Vision^45^, Triangulate^46^, scFAN^47^) is the same as the ones we enumerated for the bulk data. As we showed earlier (**Figure 1**) pyPAGE is robust to data sparsity and it accounts for biases in our annotations. As a result, we expect a reduction in the number of false positives, which makes pyPAGE especially well-suited for pathway exploration of cell-type specific gene expression patterns.

**Figure 3.**
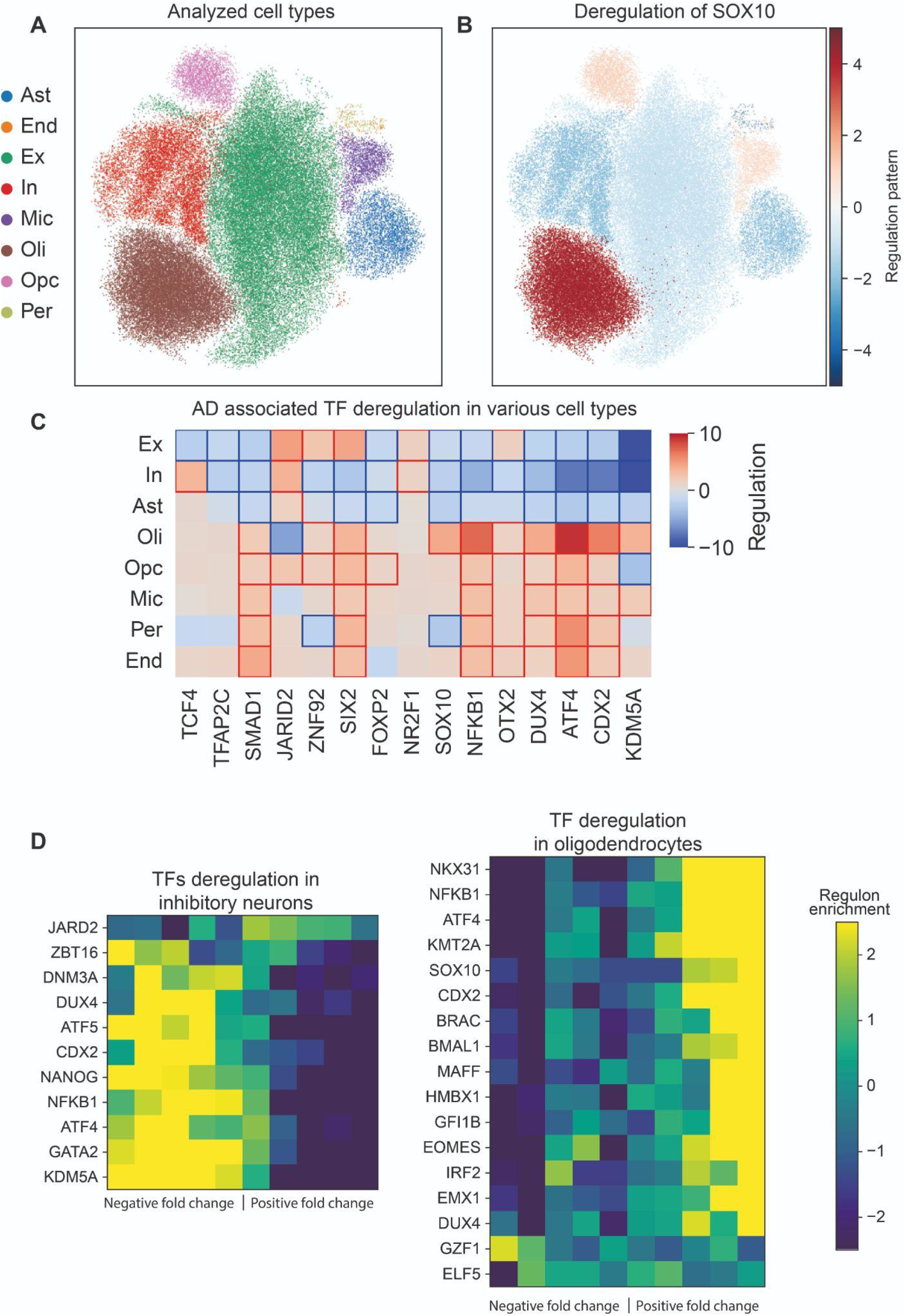
Cell type specific differential activity patterns of transcriptional factors in AD. (A) Cells from the analyzed ROSMAP dataset represented on a force-directed graph embedding. The clusters are colored according to cell-types: excitatory neurons (Ex), inhibitory neurons (In), astrocytes (Ast), oligodendrocytes (Oli), oligodendrocyte progenitor cells (Opc), microglia (Mic), endothelial cells (End), pericytes (Per). (**B)** The same cell-type clusters colored according to differential activity of SOX10 between cells from non-AD and AD samples estimated using pyPAGE. The magnitude of the regulation pattern was calculated as scaled conditional mutual information multiplied by the factor representing the direction of deregulation. (**C)** Summary of the cell-type specific deregulation patterns of the TFs identified in the analysis of the bulk data. Heatmap cells with significant associations (p-value<0.05) are framed. **(D)** Heatmap representations of concordant expression changes in expression of TF target genes in inhibitory neurons and oligodendrocytes. Here rows correspond to TFs and columns to gene bins of equal size ordered by differential expression, the cells are colored according to the enrichment of genes from regulons in a corresponding bin.

As mentioned above, the primary goal of integrating single-cell data into our AD analysis workflow was to measure contribution of the identified regulons to the gene expression of individual cell types. As expected, all the TF regulons showed significant deregulation in at least one cell type. Some of them, in fact, showed significant associations with gene expression across all major cell types, indicating their diverse roles in brain homeostasis. For example, ATF4 is known to act in a cell-type specific manner and depending on the cellular context, it activates both pro-survival and pro-death pathways^48^. Accordingly, in our analysis, we observed that theATF4 regulon showed divergent patterns of activity in various cell types. In many cases these diverging patterns reflected their underlying roles in development; for example, SOX10 is a major regulator of oligodendrocyte survival^42^ and inhibitor of astrocyte differentiation^49^ (**Figure 3B**). Or, similarly, TCF4 is known to be a major regulator of neurogenesis^50^. Finally, we observed strong evidence of epigenetic reprogramming, manifested either by activation or repression of KDM5A and JARID2 target regulons in all major cell-types. Taken together these findings illustrate cell-type specific regulatory programs involved in AD, further highlighting the generalizability of pyPAGE to single cell transcriptomic measurements.

Next, we asked whether there were patterns in the single-cell data that had remained hidden in our analysis of bulk RNA-seq. For this, we analyzed the cell-type specific differential regulon activity of factors not identified in the bulk analysis (**Figure S5, A-E and Table S3**). Interestingly, in many cell-types we observed deregulation of transcriptional programs that underlie the development of these cell types. For example, inhibitory neurons showed downregulation of factors involved in neuronal differentiation (DMT3A^51^, NFKB1^52^, KDM5A^53^) (**Figure 3D, left panel**). Likewise, in excitatory neurons we observed inhibition of TF regulons which characterize mature neurons (NCOR2^54^, KDM5A^53^) and activation of regulons controlled by protective factors (SIX2^55^, NR1H3^56^, TCF7^57^) (**Figure S5B**). Glial cells showed general upregulated developmental and cytoprotective factors, demonstrated by activation of regeneration programs in oligodendrocytes (NKX3-1^58^, NFKB1^59^, EMX1^60^, IRF2^61^) (**Figure 3D, right panel**). Together, these observations support the hypothesis that AD pathology disrupts gene expression programs that give rise to and maintain cellular identity in neurons^62^, while simultaneously triggering cytoprotective response in glia.

### Systematic identification of RNA-binding proteins associated with pathological post-transcriptional gene regulation in AD

In the previous sections, we explored AD-associated transcriptional programs and their influence on gene expression in AD pathology. Here we aim to investigate the role of post-transcriptional regulators on gene expression changes. Many aspects of post-transcriptional regulation cannot be captured by RNA sequencing alone, however, the RNA turnover, which is controlled by RNA-binding proteins and microRNAs, can be gleaned from RNA-seq data. To collect unbiased estimates of transcripts stability, we reprocessed MSBB RNA-seq data using our previously developed tool REMBRANDTS^25^. Then for each gene we calculated differential stability rates by comparing RNA stability modulations in two groups of samples using t-test. To predict RBP deregulation, we used an annotation of post-transcriptional regulons which we compiled and collated from multiple sources: namely ENCODE^57^ and POSTAR databases^64^, as well as the Mireya Plass study^65^. By applying pyPAGE to this newly compiled annotation of RBP targets we were able to disentangle the contribution of each RBP to the differential transcripts stability estimates and uncover novel post-transcriptional regulators associated with AD.

Utilizing the framework described above, we identified 13 candidate RBPs that are associated with differential RNA stability in AD (**Figure 4A**). Several of these proteins have been previously implicated in AD pathology. For example, in an earlier study, we had identified ZFP36 as a regulator of AD-related changes in RNA stability in an entirely independent dataset^25^. Here, pyPAGE confirmed that ZFP36 activity is indeed perturbed in AD as evidenced by upregulation of the genes targeted by this RBP. Similarly, pyPAGE identified deregulation of HNRNPK and HNRNPD, two RBPs that are silenced in AD due to epigenetic alterations^66,67^. The HNRNPK regulon is destabilized in AD samples, indicating that this RBP acts as an enhancer of RNA stability. In contrast, HNRNPC promotes the degradation of its regulon, which results in an increase in their stability in AD^66,68^. The ability of pyPAGE to capture known and novel RBP associations with AD not only confirms the utility of this approach, but also provides readily testable hypotheses around the underlying modes of regulation.

**Figure 4.**
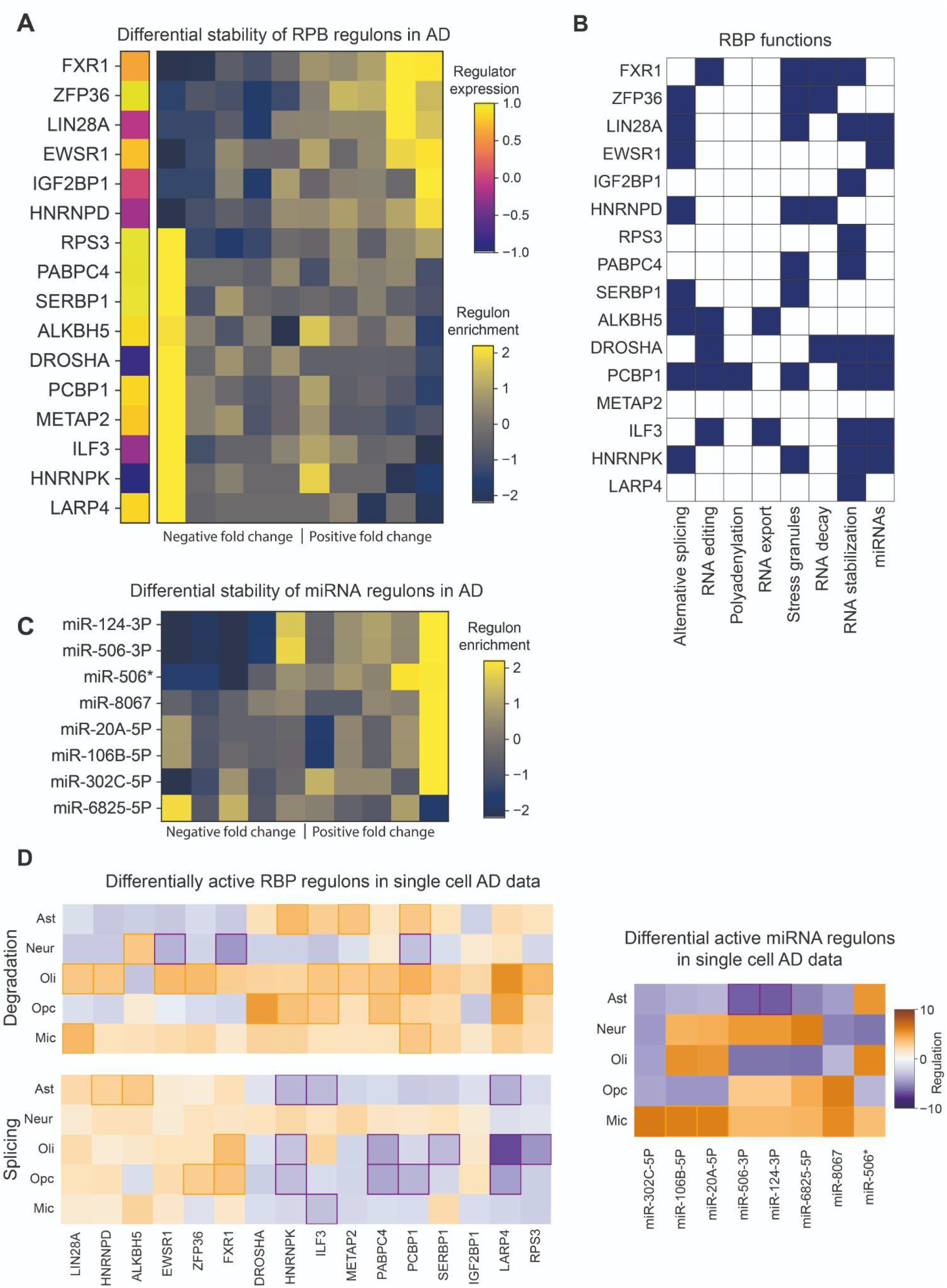
Deregulation of post-transcriptional regulatory programs in AD. (A) Heatmap representation of RBP regulons that are differentially expressed between AD and non-AD which we identified using pyPAGE. Here rows correspond to RBPs and columns to gene bins of equal size ordered by differential stability, the cells are colored according to the enrichment of genes from regulons in a corresponding bin. The leftmost column of this heatmap represents the differential expression of RBPs themselves. **(B)** Various roles performed by the identified RBPs inferred from scientific literature. **(C)** Deregulation patterns of the miRNA target gene-sets identified by pyPAGE. *miR-506 targets with GTGCCTT in their 3’ untranslated region. **(D)** Differential activity of RBP and miRNA regulons in various brain cell types. The codes for the analyzed cell-types: neurons (Neur), astrocytes (Ast), oligodendrocytes (Oli), oligodendrocyte progenitor cells (Opc), microglia (Mic). Differential activity of RBP regulons was estimated based on differential rates of RNA splicing and degradation. miRNA regulons were analyzed using only estimates of degradation rates. In these heatmaps significant associations are marked by colored frames.

To show relevance of the inferred associations in the context of AD, we explored biological functions of the identified RBPs. First, we looked at the mechanisms they use to control RNA metabolism **(Figure 4B)**. Based on existing scientific literature, we show that, for the most part, our candidate RBPs regulate RNA stability directly upon binding either through repelling or recruiting RNA degradation factors^69–77^. However, some of these RBPs employ other mechanisms. For example, HNRNPK stabilizes RNA by preventing inclusion of cryptic exons during alternative splicing which is critical in several neurodegenerative diseases^78^. Further, we performed analysis of overrepresented biological pathways within regulons of the identified RBPs to characterize their role in biological processes **(Figure S8A and Table S2)**. This analysis revealed that these RBPs play a crucial role in regulating stability of various transcriptional regulators and genes involved in molecular signaling. Differential activity of these RBPs in AD might in part explain observed deregulation of transcriptional programs via direct or indirect interactions. Additionally, the analysis of pathway enrichment within RBP regulons showed that a handful of the identified RBPs such PABC4, DROSHA, RPS3 and LARP4 are involved in the regulation of oxidative pathways which might be the root cause of AD-related mitochondrial dysfunction^79^ and metabolic deficiencies^80^.

### Identification of post-transcriptional programs governed by microRNAs in Alzheimer’s Disease

MicroRNAs are another major class of post-transcriptional regulators that have been previously associated with AD pathology^6^. Though many miRNAs have been reported to change their expression in the context of AD^82^, there has been no systematic effort to infer their activity from changes in expression of their downstream targets. We sought to fill in this gap by applying pyPAGE to analyze AD-related coordinated changes in RNA stability of all annotated sets of miRNA targets. As a result we were able to identify eight candidate miRNA regulons as associated with AD (**Figure 4C**). Overall AD-specific deregulation patterns of miRNA target genes were characterized by an increase in their stability. This suggests reduced activity of the miRNAs, which is in line with the notion that AD neurons lose the ability to process miRNAs^83^. The majority of the identified miRNAs either alter their expression in AD or are involved in several of the AD pathological mechanisms^84–88^. Functional analysis of genes regulated by these miRNAs (**Figure S8B and Table S2**) showed that they largely control neuronal processes. So, given significant deregulation patterns of the identified miRNAs, their previously described implication in AD and their role in brain functions, these miRNAs form an attractive list of targets for further focused research.

### Cell-type specific analysis of post-transcriptional regulatory programs in AD

Similar to how we described cell-type specific patterns of transcriptional regulons, we characterized the activity of RBPs and miRNAs in each of the disease-relevant cell-types. For this, we analyzed single cell RNA-seq data from Morabito et al.^20^, which contains cells from both neuronal and glial lineages. To evaluate the effects of post-transcriptional regulation in single-cell data, we used functions from the ‘velovi’ package^89^ for inference of splicing and degradation rates within each cell-type. Using non-parametric tests, we compared these transcriptomic rates between the groups of healthy and AD donors evaluating their differential values. These estimates present an unbiased view of post-transcriptional regulation by controlling for the effect of transcriptional activity. Given their known regulatory functions, we used both differential stability and splicing rates to test for differential RBP activity. For miRNAs, on the other hand, we focused primarily on differential stability rates.

This analysis confirmed that the post-transcriptional regulatory programs we had previously identified in bulk RNA-seq data were predominantly deregulated in cells of glial lineage, and to a lesser degree in neurons (**Figure 4D**). Oligodendrocytes, astrocytes and oligodendrocyte progenitors predominantly showed an increase in splicing rates and a decrease in degradation rates of RBP regulons. Neurons, on the other hand, did not exhibit broad changes in splicing rates and only a few RBP regulons showed changes in their degradation rates. Results from miRNA regulons, however, were less concordant between the single-cell and bulk data modalities. Four out of eight miRNAs showed significant deregulation patterns and only in two cell-types. We observed a significant decrease in the stability of miR-106b-5p and miR-20A-5p regulons in microglia. It should be noted that upregulation of miR-106b-5p marks microglial activation^90^. Astrocytes showed a significant increase in the stability of miR-124-3p and miR-506-3p regulons, which are similarly critical for astrocytes transdifferentiation into neurons^91^ and neuronal proliferation^92^. The key observation that most prominent patterns of miRNA and RBP regulons perturbations occur in non-neuronal cells highlights the importance of glia in AD pathology.

### pyPAGE identifies deregulation of post-transcriptional programs as a major source of heterogeneity within the cohort of Alzheimer’s patients

Finally, we employed pyPAGE for the discovery of novel sources of heterogeneity within Alzheimer’s disease patients based on the activity of our identified master regulators. To measure activity of a regulon in an individual sample, we used conditional mutual information (CMI) values that are provided by pyPAGE as part of its output. Here, the information criterion was estimated between the target genes and the standardized gene expression/transcript stability profile of each sample. As we had mentioned earlier, one of the benefits of using CMI is that it both captures monotonous changes in regulon abundance (**Figure S9A**) as well as non-linear patterns, which allows pyPAGE to efficiently measure regulon activity in individual samples.

Based on estimated activities of transcriptional and post-transcriptional regulons, we used Cox regression to find significant associations between the activity of regulons and the survival of patients. Interestingly, we found that post-transcriptional programs are most strongly associated with patient survival, represented mainly by RBP regulons (LIN28A, HNRNPD, EWSR1, FXR1, HNRNPK, ILF3, PCBP1) and a few miRNA regulons (miR-302C-5P, miR-6825-5P) (**Figure S9C**). Using activity estimates of these RBP regulons, we performed unsupervised clustering of samples. This resulted in two clusters (**Figure 5A**), showing starkly different levels of activity of several RBPs. The activity of RBPs in samples from one of these clusters was higher than those observed in healthy samples (**Figure 5C**). Remarkably, low activity of RBP regulons was associated with shorter survival, as indicated by the hazard ratio of 4.84 (p-value=0.012). In comparison, no other major biological factors had a significant effect on survival (**Figure 5D**). These results highlight the underexplored significance of post-transcriptional regulation in AD and their potential clinical utility in improving patient stratification.

**Figure 5.**
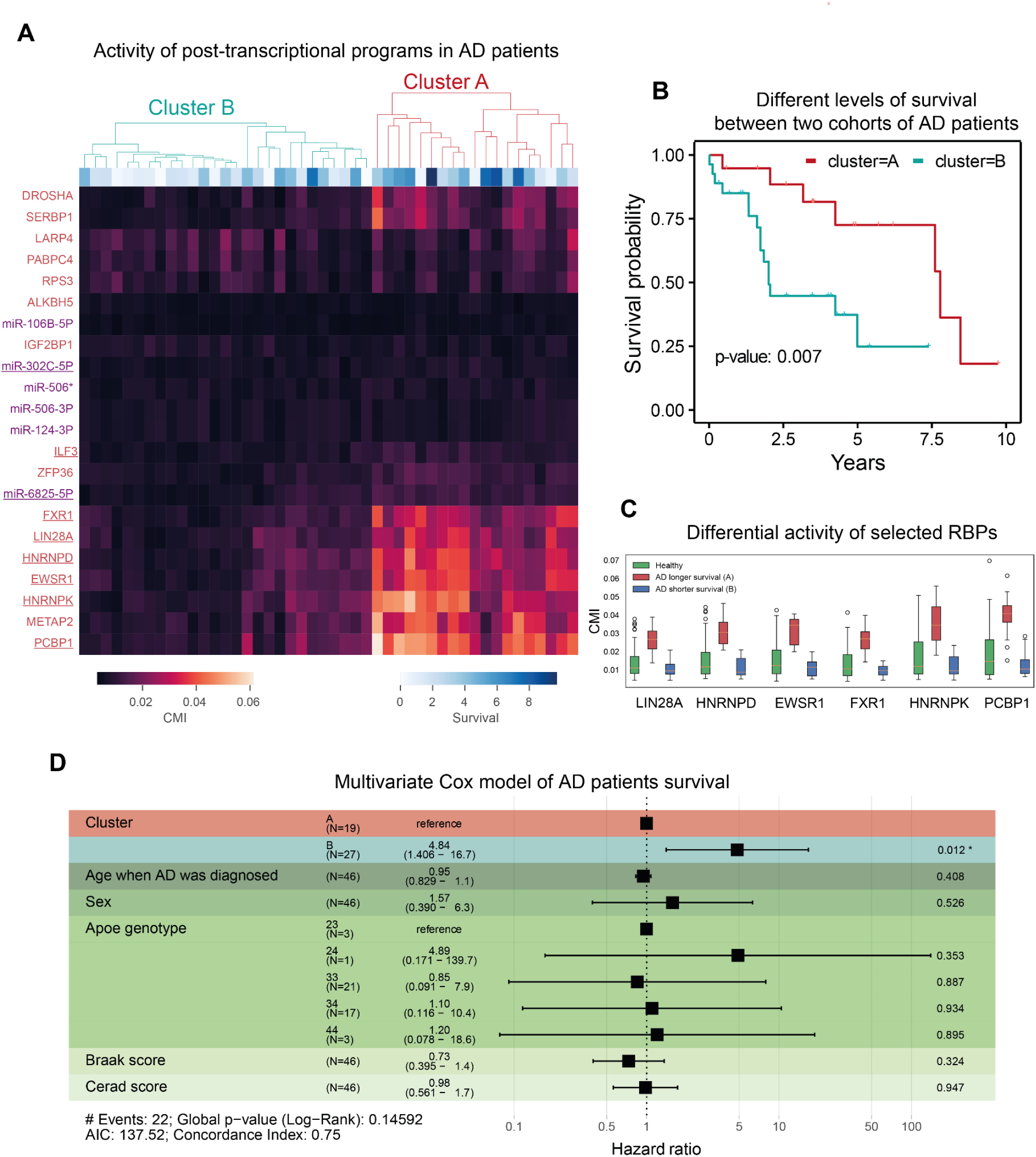
Association of activation of post-transcriptional regulation programs with survival of patients with AD. (A) Heatmap representing differences in the activity of the previously identified post-transcriptional regulons in AD samples. Factors which activity is significantly associated with survival are underscored. The dendrogram reflects the results of unsupervised clustering of samples based on activity of factors associated with survival. **(B)** Kaplan-Meier curve representing the difference in survival between two groups of patients stratified based on the activity of post-transcriptional regulons. **(C)** Comparison of the activity of selected RBP regulons in healthy samples and samples from two AD clusters. **(D)** Summary of the Cox regression analysis.

In order to identify underlying genetic root causes of the observed heterogeneity within AD patients, we performed differential analysis of genomic variants enrichment between the two identified groups of patients (**Table S4**). Specifically, we looked at the AD-related GWAS^93^ polymorphisms and polymorphisms related to genes of the analyzed RBPs. In this manner we have identified that patients with a longer lifespan are characterized by AD-related genomic variants in the upstream regions of CD55, PLK2 and SAP18 protein-coding genes. On the other hand, patients in the second group are characterized by variants upstream of STC1 and SCARA3 genes. The analysis of mutations associated with the RBPs showed that longer lifespan is associated with an upstream variant of PABPC4 gene and conversely shorter lifespan is related to two intronic variants in FXR1 and LARP4 genes. Incidentally, among these RBPs only FXR1 shows differential activity patterns between samples which makes it a primary subject for further research of the proteins which influence AD survival time.

## DISCUSSION

Understanding the patterns of gene expression regulation in a biological system is a challenging yet rewarding task which holds a potential to provide valuable insights into which factors control the behavior of a biological system. In this paper, we propose a systematic approach for exploration of gene regulation based on the analysis of regulon activity, which we implemented in our framework called pyPAGE. Two major problems which restrict the analysis of gene expression is technical variation and the combinatorial action of multiple regulatory factors. pyPAGE addresses both of these issues. First, by focusing on the differential activity of regulons rather than individual genes it becomes more robust to random variations in data. Second, by conditioning the association of gene expression with a particular regulon on a number of regulons each gene is associated with, pyPAGE effectively uses the structure of an interaction network to more efficiently disentangle perturbation responses of multiple factors. In settings with simulated data, pyPAGE performs favorably to other commonly used tools and demonstrates its robustness to noise in the input, such as, for example, data sparsity. We show that pyPAGE can be used for inference of changes in regulatory programs both on transcriptional and post-transcriptional levels. Moreover it is also able to perform analysis in the domain of single cell RNA-seq data.

In this study, we have showcased the pyPAGE by applying it to the analysis of large RNA-seq datasets, both single-cell and bulk, in the context of Alzheimer’s disease. We were able to recapitulate a number of previously known pathways and uncover new insights into the potential role of multiple TFs, RBPs and miRNAs in AD. Deregulation of the target genes controlled by these regulatory programs might explain pathological changes in AD, as they control key biological processes such as cell cycle progression, cellular signaling, neurogenesis, proteostasis and mitochondrial functions. By using single cell data of healthy and AD samples, we were able to better capture the cell-type specific patterns of the identified regulon. We found that AD-associated deregulations in gene expression in neurons are modulated primarily at the transcriptional level, whereas glial gene expression reprogramming is predominantly due to aberrant activity of post-transcriptional regulators. We further leveraged pyPAGE for the inference of regulatory programs hidden in the bulk data and uncovered characteristic upregulation of developmental TF regulons in most of the cell types. Similarly, we have identified that miR-124-3p, one of the key miRNAs that is depleted in AD brains, most strongly affects astrocytes, which might block their transdifferentiation into neurons^91^.

Like most complex human diseases, Alzheimer’s disease is a heterogeneous disease, likely with multiple molecular subtypes. Our results indicate that the activity of the various regulatory programs identified can be used to risk-stratify AD patients. In our stratification, patients with higher activity of several RBPs showed overall better clinical outcomes. It is noteworthy that these RBPs are associated with oxidative pathways. This simultaneously links activity of these RBPs to the regulation of both oxidative stress and metabolic activity. As new therapeutic strategies against AD emerge, knowledge of these regulatory programs may allow for better stratification of responders and non-responders among the patient population.

## MATERIALS AND METHODS

### Analysis of bias in gene-set annotations

To demonstrate specific properties of gene-set membership distribution, we analyzed several commonly used annotations of gene-sets. Specifically, we examined annotations of biological pathways and miRNA targets^36^ from MSigDB^3,95^, TF regulons from Vorontsov et al.^35^ and RBP regulons which we compiled ourselves. For each gene in an annotation we, first, calculated the number of gene-sets each gene is a member of, then, divided these values into several equally sized bins and then showed that this distribution can be closely approximated with the power law. In each case we report the inferred gamma parameter of the distribution and the R^2^ statistics. We observed that for several annotations the long tail of the gene-set membership distribution deviates from the power law, indicating bias in annotations.

We further explored the possible sources of bias observed in annotations. Firstly, we analyzed the association between the number of protein interactions and the protein abundance. To determine the number of interactions, we utilized the STRING database^96^ considering only physical interactions with a confidence score greater than 0.9. As a source of protein abundance data we used proteome measurements from HEK-293 cells ^97^ . Secondly, we analyzed the association between pathway membership and citation index^98^ for each gene in the annotation of biological pathways. In both cases we were able to show that there is a linear relationship between the analyzed variables indicated by high correlation values.

### pyPAGE algorithm

The computational framework implemented by pyPAGE includes three key steps. First, it preprocesses the input data by discretizing the differential gene expression profile into equally sized bins and converting a gene-set annotation into a binary format. Second, it performs estimation of conditional mutual information between differential gene expression and each gene-set. Here the metric is conditioned on the gene-set membership profile which is also discretized into several bins. Third, the program estimates the significance of the inferred associations. For this, a non-parametric testing procedure is implemented. The differential expression profile is randomly shuffled multiple times and at each iteration the conditional mutual information is calculated between these profiles and the scrutinized gene-set. Then the significance of the association is inferred as the proportion of cases when the CMI of randomly permuted differential expression profile turned out to be greater than CMI of the original profile. This procedure is applied to a list of gene-set in a descending order of the CMI calculated at the second step. When multiple insignificant associations (by default 5) come in a row, the testing stops. pyPAGE may also use an additional filtering procedure which is applied from top-to-bottom to the list of most informative gene-sets. Each next gene-set is required to fulfill

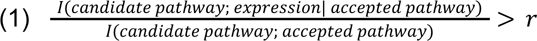

In the end pyPAGE presents the results in the form of a table. It also has an option of the graphical output, which is a heatmap representing gene-set members enrichment within equally sized bins ordered by differential gene expression.

pyPAGE is implemented as a python package and is openly available at https://github.com/goodarzilab/pypage.

### Tests

In order to validate various aspects of pyPAGE execution we have developed a series of tests based on simulated data. Our main focus was on proving that the program is robust to gene-set annotation biases. To do that we evaluated its average performance using artificial datasets and benchmarked it against other programs. We also conducted two additional tests to demonstrate consistency of pyPAGE predictions. Though to produce highest performance pyPAGE needs to receive clean data with strongly manifesting biological signals, we still expected the program to perform decently in various adversarial cases. By subsampling the original input and by modulating the robustness of signals encoded in the data, we wanted to show at which cost the reduction in input quality is to the overall performance. The full description of the tests is provided further.

### Simulating differential expression

All implemented tests required simulating differential gene expression data. Though in principle pyPAGE does not depend on any particular type of the input distribution, we decided to make the input data realistic. So, we based the simulation on the actual gene expression estimates from a real biological system, which in this case was activation of murine CD4 T cells (PRJNA296380)^99^. Starting off with fastq files, we first applied Salmon^67^ to quantify gene counts and afterwards applied DESeq2^68^ to estimate differential gene expression between two groups of samples. To produce an arbitrary amount of simulated gene expression profiles using this data, we performed random shuffling of the data across genes. This procedure minimized the informational content which had been in the data initially. We demonstrated that by performing pyPAGE gene-set enrichment analysis of terms from biological pathway annotation using original and augmented data. While the analysis of the original data reported 76 pathways, using simulated data pyPAGE on average reported only 3.36 pathways (standard deviation=1.80). Though this procedure removes almost completely any informational content, it preserves the properties of the original data distribution. Moreover it can be repeated multiple times to produce many samples of simulated differential expression profiles.

### Simulating pathway annotations

In addition to the differential expression profiles, each test needed a simulated pathway annotation. To be able to assess the performance of pyPAGE at later stages, during annotation design whether a gene-set is deregulated or not should be directly encoded in its structure. And so our goal was to create annotations each containing in total 1100 gene-sets, among which 100 are known to be deregulated in the corresponding differential gene-expression profile. The number of genes that went into a newly designed gene-set chosen based on the distribution of sizes of TF regulons^35^. To simulate unperturbed gene-sets we randomly picked the necessary number of genes. To imitate deregulated gene-sets, we used genes that are either over- or under-expressed in the corresponding simulated data; the details of simulating such gene-sets are described in the following paragraph.

When simulating deregulation our objective was to imitate patterns observed in real systems, so as to avoid making the task of predicting gene-set deregulation too easy or too difficult. In a real case scenario genes of a deregulated gene-set are not always exclusively found among the most overexpressed or underexpressed genes, rather the signal is distributed over the whole range of values. Likewise, in our simulation we built probabilistic models to control which genes go into a newly formed gene-set. We aimed that the resulting distribution of differential gene expression in each gene-set should match the truncated normal distribution on an interval between -1 and 1 with predefined parameters. So for each gene-set we first defined a distribution of expected differential gene expression, then randomly sampled a number of values from it and selected those genes from a simulated differential expression profile which most closely matched these values. Parameters of the expected distribution were also selected randomly. For an upregulated gene-set, its mean was picked randomly from a random distribution in an interval between 0.5 and 1, for a downregulated gene-set the interval was between -1 and -0.5. The standard deviation was uniformly selected in the range between 0.1 and 0.2. To simulate dual regulation, we mixed genes selected for an upregulated gene-set with those of a deregulated one. Finally, to imitate biological and technical noise to each gene-set we added a randomly selected group of genes. The ratio of randomly added genes was controlled by signal-to-noise ratio, which in the general case was set to 33%.

The resulting gene-set annotations were unbiased with gene-set membership following approximately normal distribution. To produce biased annotations, we needed to add bias externally. And so we selected 2000 genes which would represent genes with promiscuous activity. Then each gene-set in an annotation was concatenated with additional 100 genes randomly selected from this group. The idea behind this approach is that added genes would occur much more frequently than all other genes effectively creating bias in gene-set membership distribution.

### Benchmarking pyPAGE

Using the artificial datasets containing simulated differential expression and gene-set annotations, we were able to analyze pyPAGE effectiveness and compare it with other programs: iPAGE and fgsea. The test was basically a classification task: each program needed to predict deregulated gene-sets. The performance was judged based on the knowledge of true patterns of regulation which were encoded in simulated annotation. We used multiple metrics to assess performance of programs which were averaged across 20 iterations with different sets of simulated data. To present these results we plotted ROC (receiver operating characteristic) curves, PR (precision recall) curves, DET (detection error tradeoff) curves, FDR (false discovery rate curves) curves and calculated respective AUC (area under the curve) scores. Since the number of unperturbed gene-sets was much greater the most relevant metric in this task was PR-AUC score. Programs were tested in 4 different conditions to reflect the effect of bias and non-linear regulation on the quality of predictions.

### Robustness test

Since in practical applications, the input data of a gene-set can often be corrupted, we developed two tests to explore how robust pyPAGE is to the augmentation of the input. The first strategy that we implemented for simulating input perturbation was increasing the amount of noise in the data. In the previous we set signal-to-noise ratio to 33%, however here we varied this parameter in the range between 0 and 100%. The second type of augmentation we used was implemented by randomly subsampling the input data. We iteratively decreased the number of genes in the input by 5% measuring pyPAGE performance. Both tests output pyPAGE sensitivity changes in response to input modulation.

### Analysis of Alzheimer’s Disease

To characterize perturbation of transcriptional and post-transcriptional programs in AD, we analyzed gene expression in 197 samples of BM36 brain regions from MSBB study^18^. Specifically, we narrowed down analysis to comparison between the two groups of donors: those without pathology and those with definite pathology. To analyze changes in gene expression, we processed the input fastq files with Salmon^100^ and then applied DESeq2^22^. The analysis of clustering of samples based on their gene expression showed 4 large clusters which corresponded to the batch group and sex of the donor. Within each cluster there were both healthy and pathological samples. However, since clusters corresponding to the second batch group showed poor separation between samples, they were excluded from the analysis (in total 83 samples were removed). Within each of the two remaining clusters corresponding to female and male samples we estimated logarithms of fold changes and p-values for each gene. In order to account for any sex-specific variation, we combined the estimated p-values using a modified inverse chi-square method for correlated tests^101^ and computed the mean between logarithms of fold changes.

Afterwards we quantified differential gene expression using the following formula:

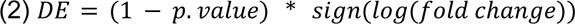

Using these estimates, we performed pyPAGE analysis of TF regulons from Vorontsov et al.^35^. In order to analyze post-transcriptional regulation, we first needed to estimate differential transcripts stability. To do this we first applied Salmon^100^ to quantify separately exonic and intronic counts and then applied REMBRANDTS^25^ to produce unbiased estimates of stability for each sample. Next using t-test we compared stability between pathological and normal samples; the resulting p-values and fold changes were used to quantify differential stability similarly to how we estimated differential gene expression. Finally, based on these estimates pyPAGE was able to infer deregulation patterns of RBP regulons from our newly compiled annotation and microRNA targets^36^ from MSigDB^3,95^.

Cell-type specific deregulation patterns were estimated using single cell data. For this purpose we used two datasets. One was an already processed dataset presented by ROSMAP study^19^. We used it for the analysis of TF regulons. There within each cellular type cluster we applied t-test to estimate differential gene expression between cells from AD and non-AD samples. Then pyPAGE was used to estimate deregulation of TF regulons within each cell type. For the analysis of RBP and miRNA deregulation, we used another single cell RNA-seq dataset from Morabito et al.^20^. We processed this dataset de novo using cellranger^102^ and velocyto^103^ in order to finally run velovi^89^. We applied velovi to single cell datasets of individual samples in order to estimate patient specific rates of RNA splicing and degradation in each cellular type. In the end in a manner similar to how we estimated differential gene expression, here we used t-test to estimate differential kinetic rates between patients with and without AD. Based on the differential splicing and degradation rates, using pyPAGE we were able to infer the activity of RBP regulons. For the analysis of miRNA regulons, we used only differential degradation rates.

We also analyzed the association between the activity of the identified regulons and patients’ survival. To do this, we analyzed another bulk RNA-seq dataset from the ROSMAP study^104^ stored in the Synapse database under syn21589959. The samples were from dorsolateral prefrontal cortex and posterior cingulate cortex. We considered only samples from AD patients for whom the age the disease was first diagnosed was not censored. This left us with 46 samples. For these samples we performed gene expression quantification using salmon and then normalized these estimates. We then used this normalized gene expression to quantify activity of TF regulons in each sample using pyPAGE estimates of conditional mutual information. Similarly, we used salmon and REMBRANDTS to estimate transcripts stability in each sample and afterwards used normalized stability to infer activity of RBP and miRNA regulons. Next, we used regulon activity in Cox Proportional-Hazards Model to identify factors associated with survival of AD patients. Based on this we selected regulons which were significantly associated with survival. We then analyzed patterns of activity of these regulons in various samples and effectively excluded those regulons which did not show any marked difference in their activity between different samples. Then based on the activity of the remaining regulons we performed clustering of samples which resulted in two big clusters. Next, we used this stratification to analyze its association with survival. For this, we built Kaplan-Meier curve and performed Cox Proportional-Hazards Model which incorporated other AD relevant covariates such as sex, APOE genotype, Braak score, Cerad score.

In order to identify traits which characterize the samples on a genetic level, we analyzed genomic variants data from the ROSMAP study. We considered polymorphisms that are associated with AD that are available in GWAS catalog^93^ and those that are associated with the analyzed RBPs. Specifically, we filtered variants for those that are enriched in one group of samples (present in over 80% of samples) or depleted in another (present in fewer than 20% of samples).

### Characterization of regulon functions

We describe the functions of the identified TF, RBP and miRNA regulons based on the enrichment of molecular pathways among their members. To do that, we used a statistical overrepresentation test from PANTHER^105^. Then we aggregated the inferred functions of each regulon and presented them in a heatmap form.

For RBPs we also characterize their roles in various RNA metabolic processes. This was done based on manual curation of scientific literature. We describe RBP associations with following molecular mechanisms: alternative splicing^106–113^, RNA editing^114–116^, polyadenylation^117^, RNA export^118,119^, stress granule formation^120,121,121–126^, RNA-decay^68,127–129^, RNA stabilization^69–77^ and miRNA functions^129–134^.

### ENCODE processing

The ENCODE eCLIP data^63^ was available for two cell lines: HepG2 and K562. In total, there were 234 <RBP, cell type> pairs (687 samples including replicates, and matched controls for each RBP). RBPs with no matched control or with matched control only (without the experiments) were excluded from the analysis resulting in the final set of 222 <RBP, cell type> pairs. The data were analyzed as follows: (1) reads were preprocessed as in the original eCLIP pipeline^135^, (2) trimmed reads were mapped to the hg38 genome assembly using hisat2^136^, (3) the aligned reads were deduplicated^135^, (4) the uniquely and correctly paired reads were filtered with samtools^137^, (5) gene-level read counts in exons were obtained with plastid^138^ using gencode v31 comprehensive gene annotation^139^, therefore joint read counts were obtained for genes with overlapping annotations, (6) differential expression analysis against matched controls was performed with edgeR^24^, (7) Z-Scores were estimated for each gene across data for all RBPs for the fold change values estimated in (6). Based on (6) and (7) reliable RNA targets of each RBP were defined as those passing 5% FDR, raw log2FC > 0.5, Z-Score(log2FC) > 1.645. The processed data is stored in HepG2_gencode_v31.tsv and K562_gencode_v31.tsv supplementary files.

## Supporting information

HepG2 eCLIP data

K562 eCLIP data

## ACKNOWLEDGEMENTS

We thank Andrey Buyan and Ivan V. Kulakovskiy for providing reprocessed ENCODE eCLIP data.

## Notes

### Competing Interest Statement

The authors have declared no competing interest.

