## Supplementary Information for "Addressing biases in gene-set enrichment analysis: a case study of Alzheimer’s Disease"

### Inventory

**Figure S1.**
**Figure S2.**
**Figure S3.**
**Figure S4.**
**Figure S5.**
**Figure S6.**
**Figure S7.**
**Figure S8.**
**Figure S9.**

**Table S1.**
**Table S3.**
**Table S4.**

### SUPPLEMENTARY FIGURES

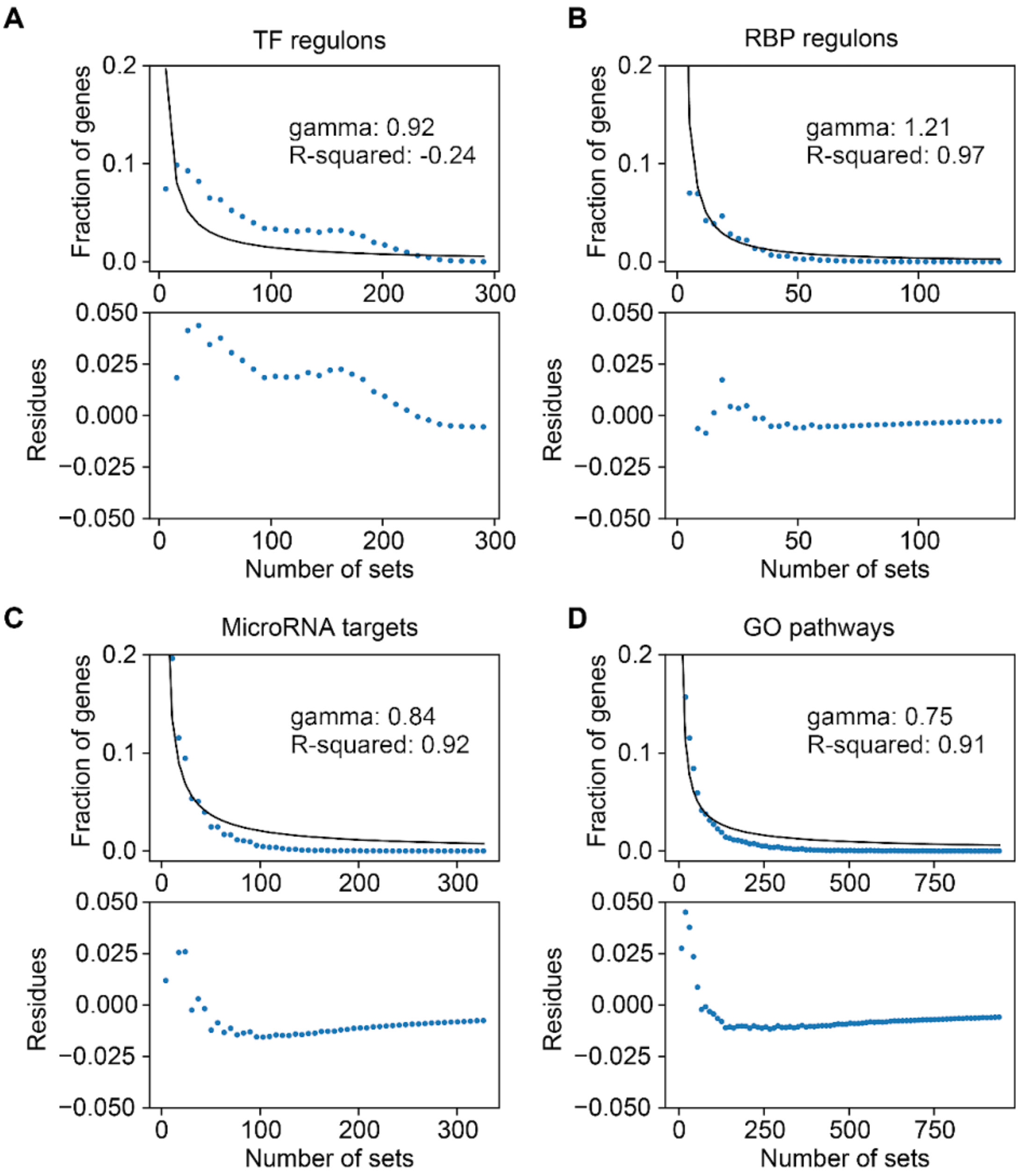


**Figure S1. Bias in gene-set annotations. (A)** Characteristics of a gene-set membership degree distribution in the TF regulons annotation. The top plot represents the observed distribution with a power law function being fit to it. We also report the gamma parameter of this distribution and the $R^2$ The bottom plot represents the deviation of the observed distribution from the power law. **(B)** Similar representation for RBP regulons. **(C)** Similar representation for miRNA targets. **(D)** Similar representation for GO pathways.

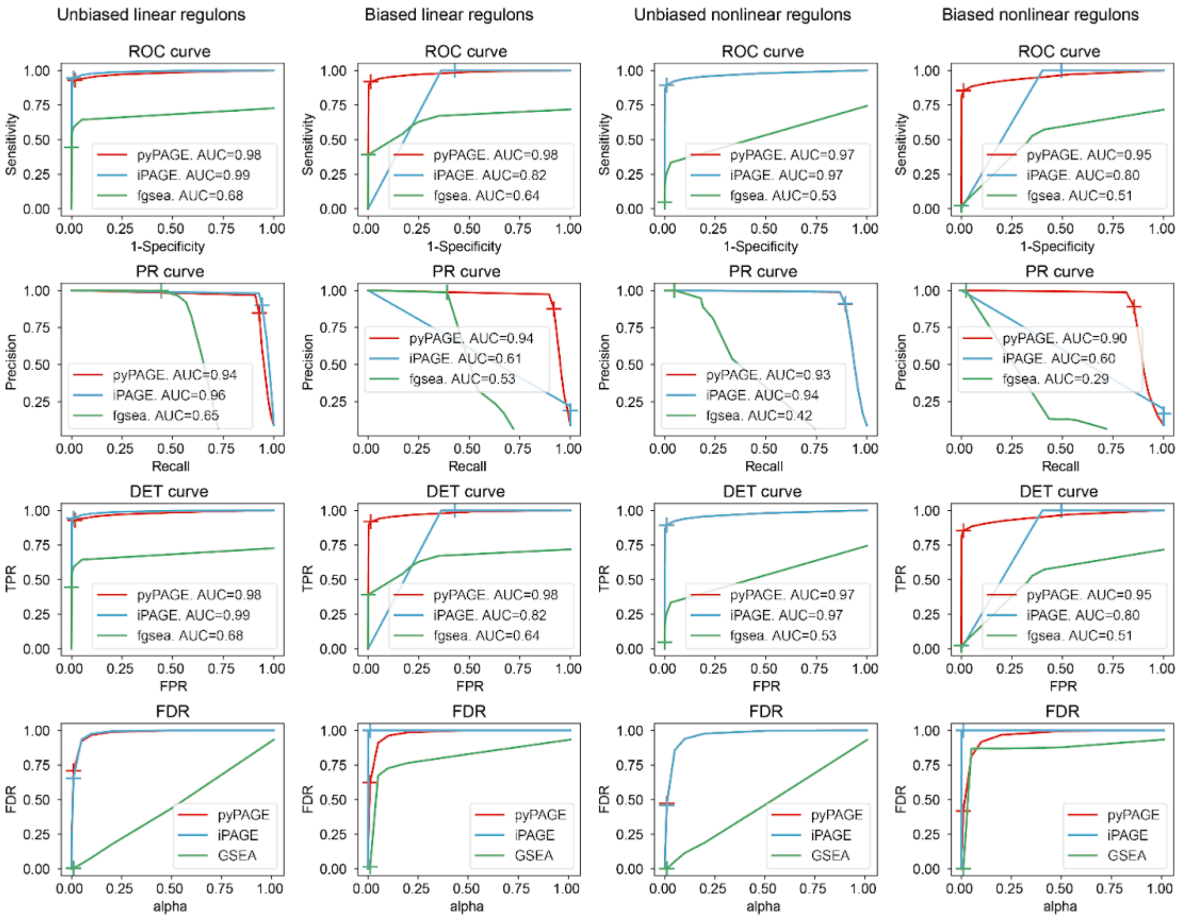


**Figure S2.**
Performance of pyPAGE benchmarked against iPAGE and fgsea in various simulated conditions. For comparison we used multiple metrics, results are presented as ROC, PR, DET curves and plot of dependency between alpha and FDR.

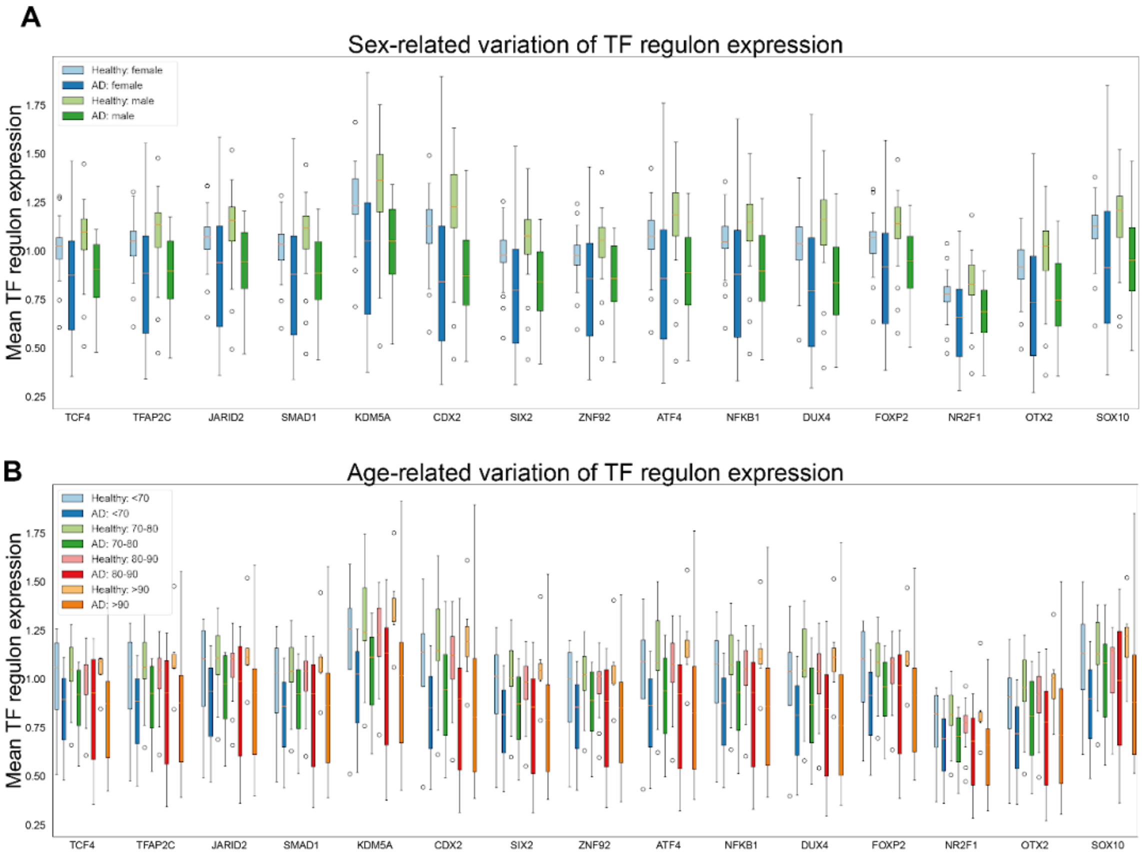


**Figure S3. Variation in expression of TF regulons.**

**(A)** Boxplots representing expression of TF regulons which were identified using pyPAGE in AD and non-AD samples from female and male donors. **(B)** Boxplots representing expression of the same TF regulons in AD and non-AD samples in different age cohorts.

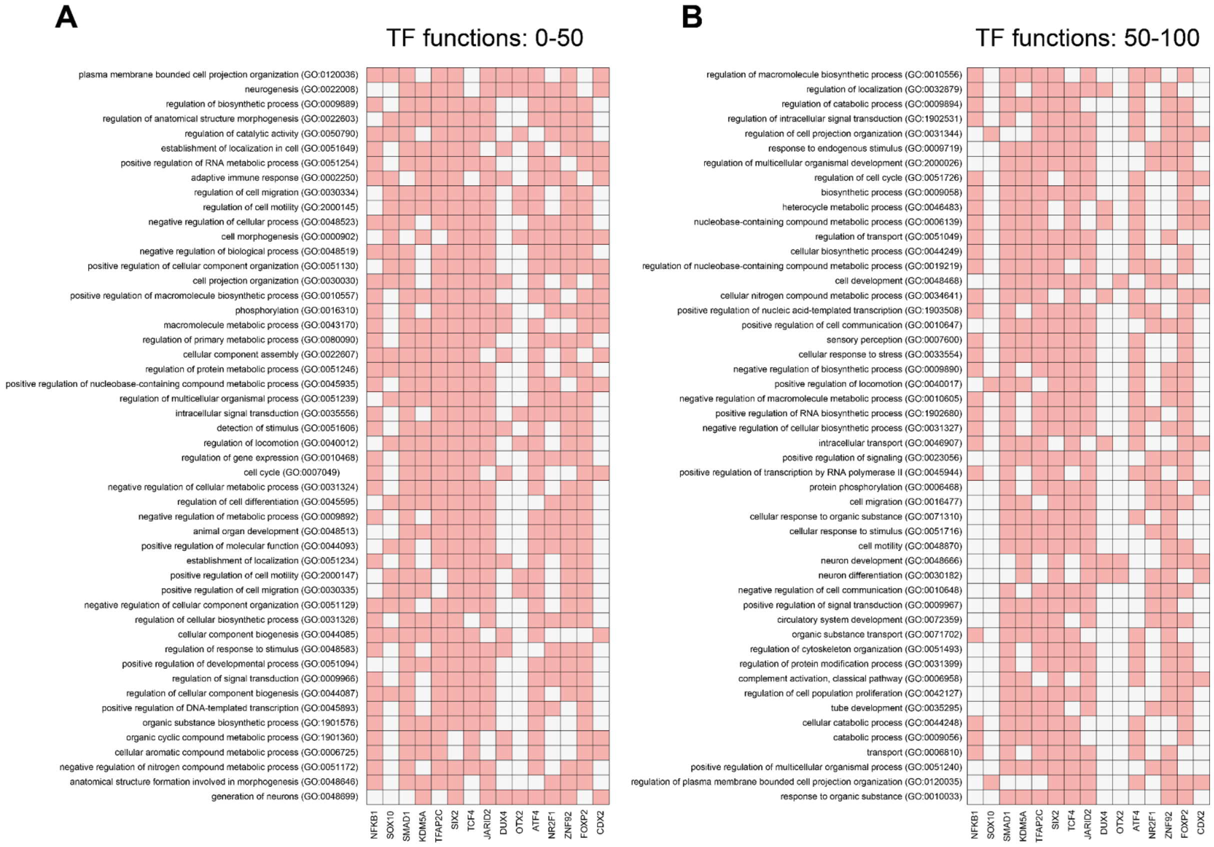


**Figure S4. Top 100 of pathways enriched in TF regulons.**

**(A)** Heatmap representing biological pathways that are enriched among targets of TFs we have identified in our analysis. Here we present top 50 pathways which are significantly enriched in most of the regulons excluding several generic terms, like “molecular process”. **(B)** This heatmap represents another 50 pathways significantly associated with the analyzed TFs.

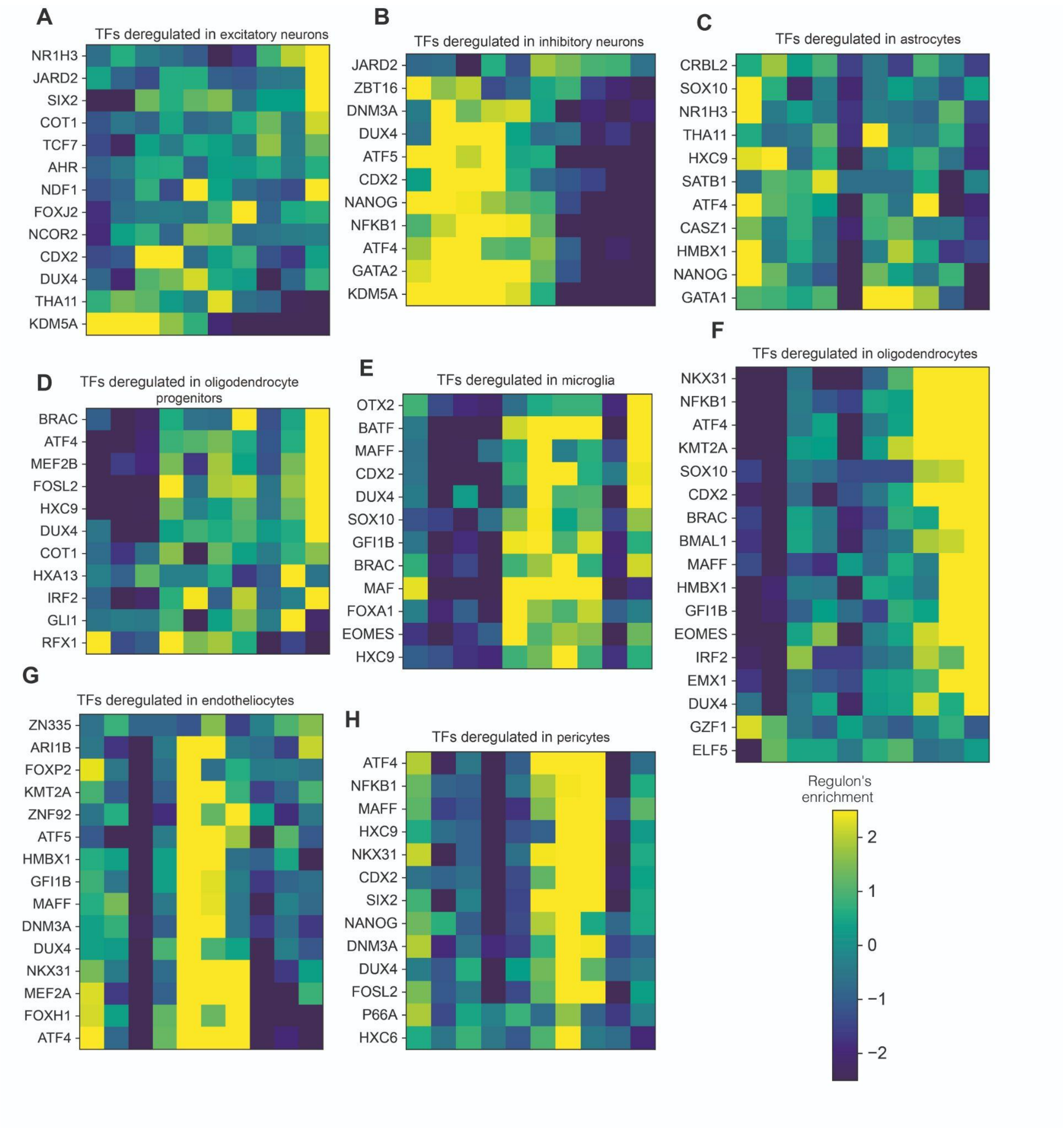


**Figure S5. Cell-type specific patterns of TF target genes deregulation in AD.**
**(A)** Differential activity of TFs in excitatory neurons inferred using pyPAGE based on the analysis of single-cell RNA-seq data. **(B)** TF regulons deregulated in inhibitory neurons. **(C)** TF regulons deregulated in astrocytes. **(D)** TF regulons deregulated in oligodendrocyte progenitors. **(E)** TF regulons deregulated in microglia. **(F)** TF regulons deregulated in oligodendrocytes. **(G)** TF regulons deregulated in endotheliocytes. **(H)** TF regulons deregulated in pericytes.

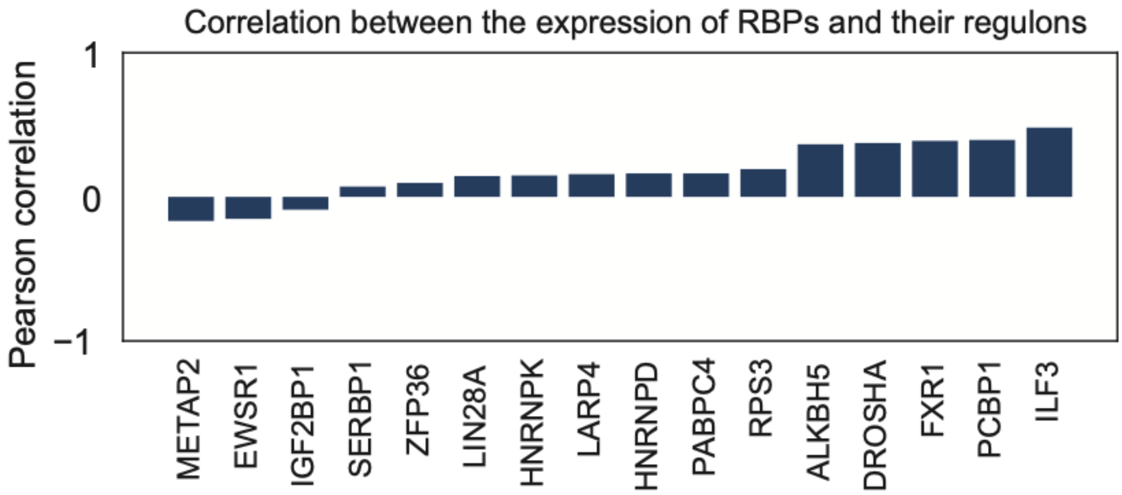


**Figure S6. Correlation between the expression of RBPs and their regulons' average stability.** The barplot representing the correlations between the expression of RBPs and the average stability of their regulons.

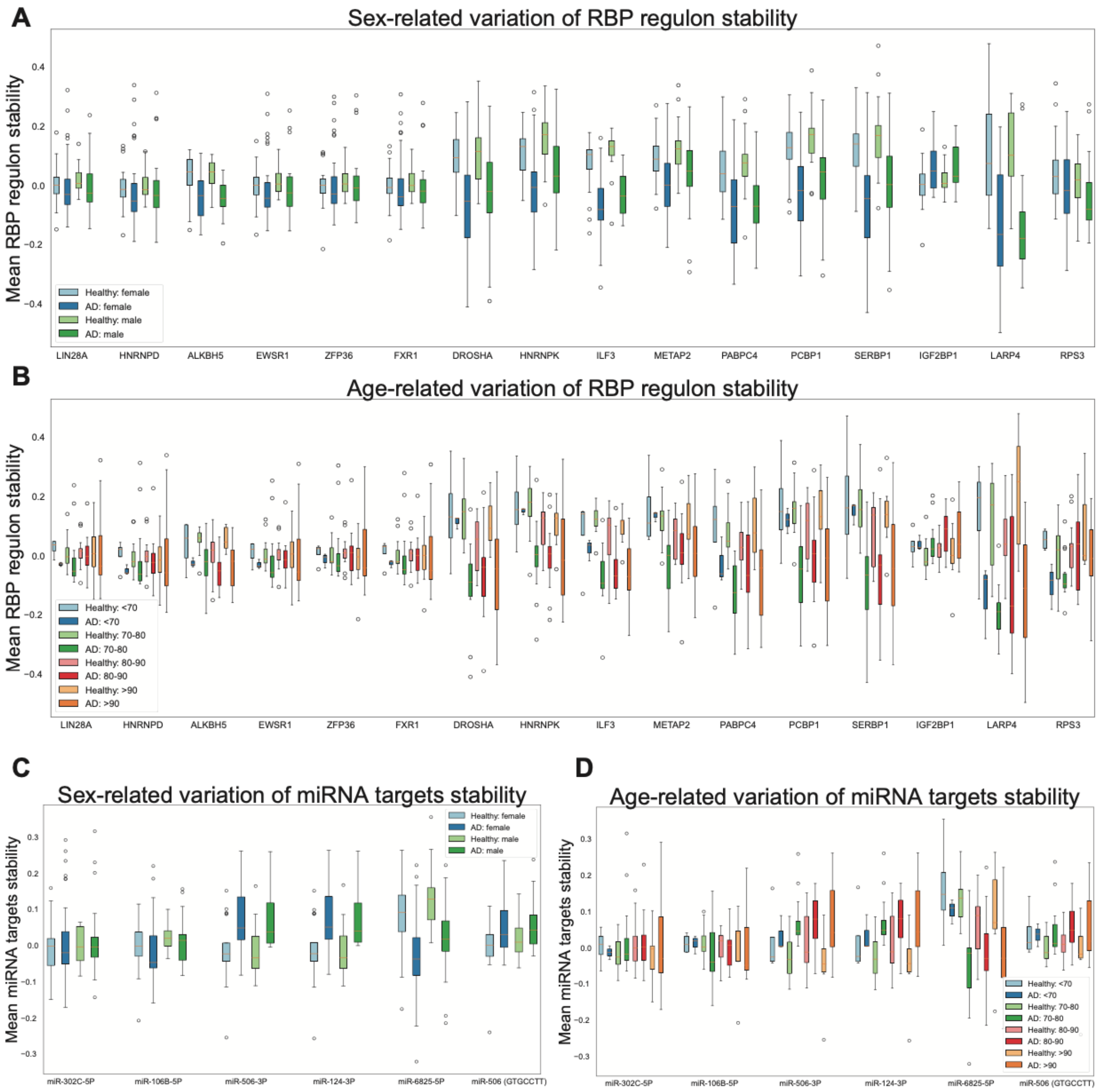


**Figure S7. Variation in expression of TF regulons.**

**(A)** Boxplots representing stability of RBP regulons which were identified using pyPAGE in AD and non-AD samples from female and male donors. **(B)** Boxplots representing stability of the same RBP regulons in AD and non-AD samples in different age cohorts. **(C)** Boxplots representing stability of miRNA targets within AD and non-AD samples from female and male donors. **(D)** Boxplots representing stability of miRNA targets within AD and non-AD samples from different age cohorts.

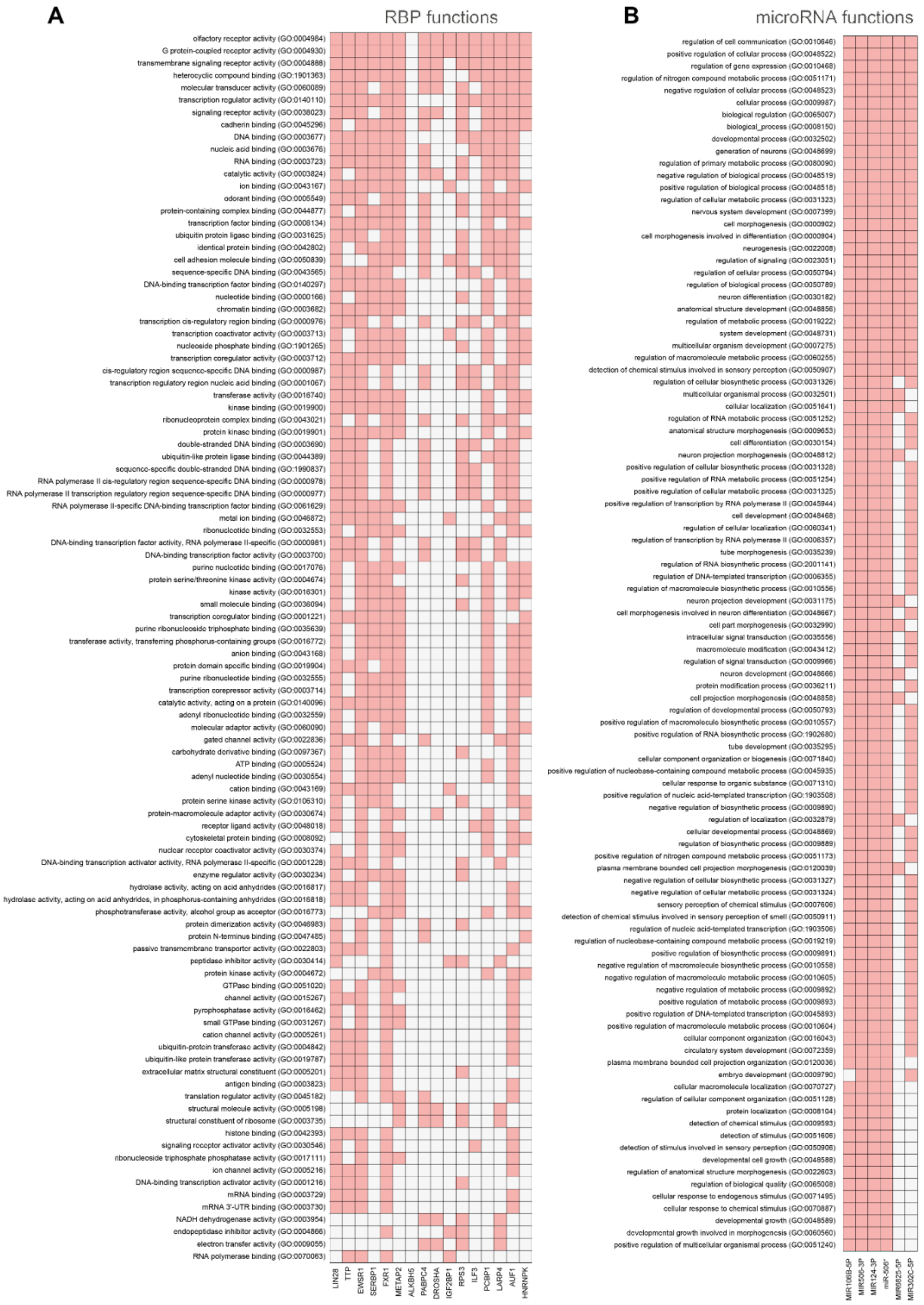


**Figure S8. Top 100 pathways enriched in RBP and miRNA regulons.**
**(A)** Heatmap representing biological pathways that are enriched among targets of TFs we have identified in our analysis. Here we present only the top 100 pathways which are significantly enriched in most of the regulons. **(B)** Similar representation for miRNA targets.

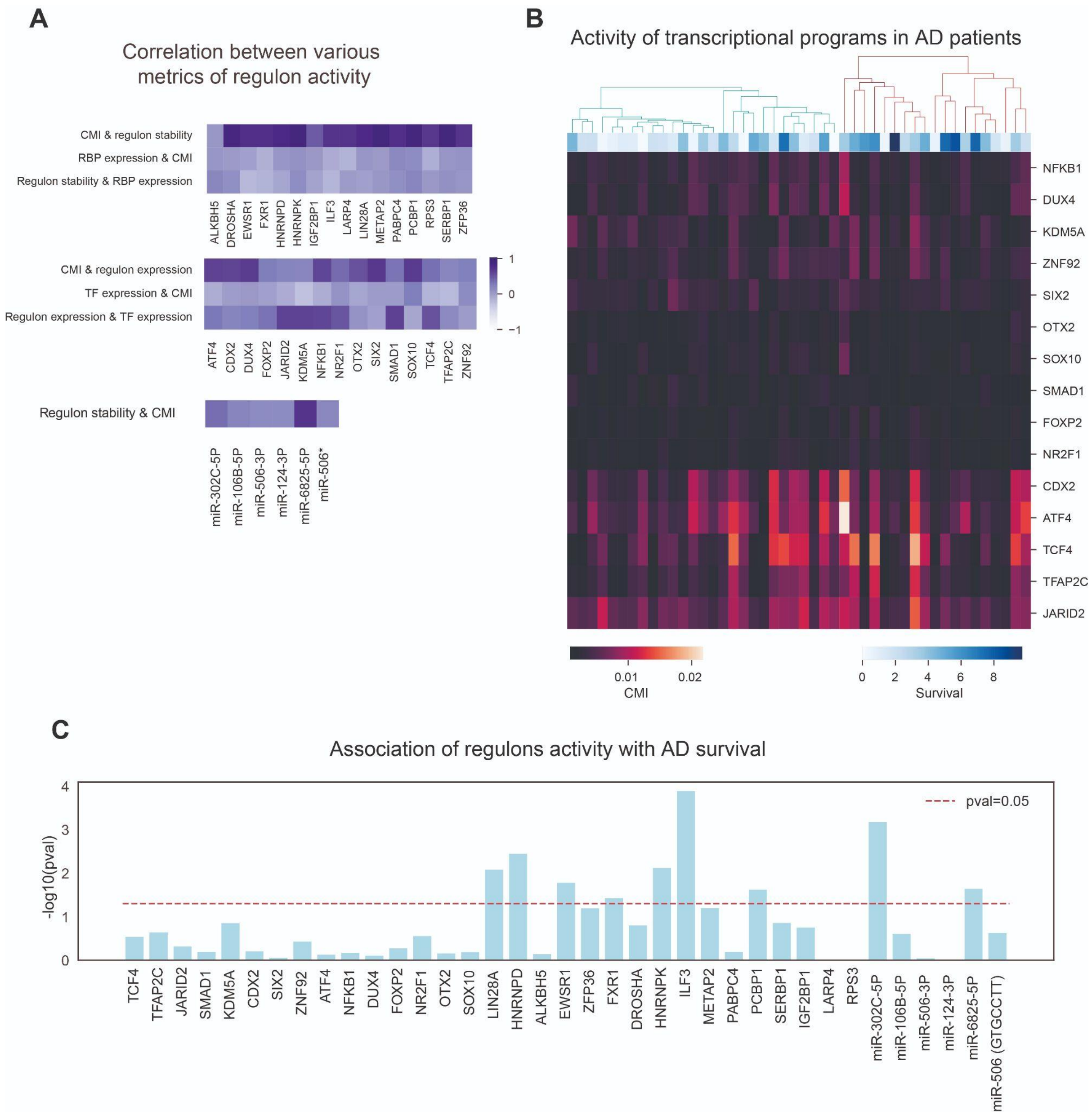


**Figure S9. Association of expression of a subset of HOX genes with survival of patients with AD**

**A** Comparison of the similarity between various regulon activity metrics, namely conditional mutual information (CMI), mean regulon abundance and the expression of a factor itself. **B** Heatmap representation of activity of transcriptional regulons in different patients. **C** Bar Plot representing significance of association of activity of each previously identified regulon with patient’s survival.

### SUPPLEMENTARY TABLES

**Table S1. Summary of pyPAGE findings**

| | |
|---|---|
| **Transcriptional factors** | NR2F1, ATF4, TFAP2, CDUX4, SOX10, ZNF92, NFKB1, KDM5A, JARID2, FOXP2, OTX2, CDX2, SIX2, SMAD1, TCF4 |
| **RNA binding proteins** | ILF3, LIN28A, DROSHA, METAP2, PABPC4, ZFP36, HNRNPK, HNRNPD, LARP4, SERBP1, RPS3, ALKBH5, IGF2BP1, PCBP1, EWSR1, FXR1 |
| **miRNAs** | miR-106B-5P, miR-20A-5P, miR-124-3P, miR-6825-5P, miR-506-3P, miR-8067, miR-302C-5P, miR-506 (GTGCCTT) |

**Table S3. Summary of pyPAGE single-cell analysis of transcriptional deregulation**

| | |
|---|---|
| **Excitory neurons** | KDM5A, SIX2, JARID2, NR1H3, CDX2, NCOR2, NR2F1, TCF7, DUX4, NEUROD1, FOXJ2, AHR, THA11 |
| **Inhibitory neurons** | KDM5A, CDX2, ATF4, NFKB1, GATA2, JARID2, DUX4, ATF5, NANOG, DNM3A, ZBT16 |
| **Oligodendrocytes** | NKX3-1, ATF4, NFKB1, KMT2A, CDX2, SOX10, MAFF, BRAC, EOMES, GFI1B, HMBX1, EMX1, DUX4, ELF5, IRF2, GZF1, ARNTL |
| **Oligodendrocyte progenitor cells** | BRAC, FOSL2, HOXC9, ATF4, RFX1, MEF2B, DUX4, IRF2, GLI1, NR2F1, HXA13 |
| **Astrocytes** | ATF4, NANOG, GATA1, NR1H3, SOX10, CRBL2, CASZ1, HMBX1, HOXC9, SATB1, THA11 |
| **Microglia** | BATF, MAF, CDX2, BRAC, DUX4, HOXC9, OTX2, SOX10, GFI1B, MAFF, FOXA1, EOMES |
| **Pericytes** | ATF4, SIX2, NKX3-1, FOSL2, CDX2, HOXC9, NFKB1, MAFF, DUX4, NANOG, HXC6, DNM3A, GATAD2A |
| **Endotheliocytes** | ATF4, MEF2A, NKX3-1, FOXH1, ATF5, DUX4, KMT2A, HMBX1, ARID1B, DNM3A, MAFF, ZNF92, ZN335, GFI1B, FOXP2 |

**Table S4. Differential enrichment of genomic variants in clusters identified based on RBP activity**

| | | Position (hg37) | Associated genes |
|---|---|---|---|
| AD associated variants | Enriched in cluster A | 10:130346154 | LINC02667 (upstream) |
| | | 1:207457845 | CD55 (upstream) |
| | | 5:57600044 | PLK2 (upstream) |
| | | 13:21712984 | SAP18 (upstream) |
| | | 14:29629456 | RNU11-5P, RNU6-864P (upstream) |
| | Enriched in cluster B | 12:59221812 | LINC02388 (upstream) |
| | | 8:23571807 | STC1 (upstream) |
| | | 8:27487790 | SCARA3 (upstream) |
| | | 8:142611971 | - |
| RBP associated variants | Enriched in cluster A | 1:40045083 | PABC4 (upstream) |
| | Enriched in cluster B | 3:180643165 | FXR1 (intron variant) |
| | | 12:50820467 | LARP4 (intron variant) |
| | | 12:50820468 | LARP4 (intron variant) |
